# The metaplastic precursor state to oesophageal adenocarcinoma represents reversion to a transient epithelial cell state in the developing oesophagus

**DOI:** 10.1101/2024.07.25.605105

**Authors:** Syed Murtuza Baker, Aoibheann Mullan, Rachel E Jennings, Karen Piper Hanley, Yeng Ang, Claire Palles, Neil A. Hanley, Andrew D. Sharrocks

## Abstract

In Barrett’s oesophagus (BO), the precursor of oesophageal adenocarcinoma, the adult stratified squamous epithelium is replaced by a simple columnar phenotype. This has been considered metaplasia; the inappropriate conversion from one adult cell-type to another. In fact, BO could be a reversal of mammalian embryogenesis when the early foregut is first lined by simple columnar epithelium. Exploring this hypothesis has been hampered by inadequate molecular details of human oesophageal development. Here, we adopted single cell transcriptomic and epigenomic approaches to discover and decode the cell types that constitute the initial primitive columnar, transitory and subsequently stratified lower oesophageal epithelium. Each stage is comprised of several previously undefined epithelial sub-populations. HNF4A, a major driver of the BO phenotype, is a prominent transcriptional regulator in the early foregut columnar cells, but not in the later ciliated or stratified cells, and is central to gene regulatory programmes known to be reactivated in BO. Moreover, GWAS susceptibility SNPs for BO mapped to putative regulatory regions in fetal epithelial cells, which are inaccessible in the corresponding adult epithelial cells. Collectively, these data argue that the path to BO involves reactivation of pathways that define primitive fetal epithelial cell states.

## INTRODUCTION

Oesophageal adenocarcinoma (OAC) carries major mortality and morbidity, with a 5-year survival less than 20%. Incidence has risen markedly over recent decades and has been linked strongly to a background of metaplasia, adult-to-adult cell conversion, in the lower third of the oesophagus associated with gastro-oesophageal reflux disease (Smyth et al., 2017). This metaplasia is termed Barrett’s oesophagus (BO) where the normal multi-layered (stratified) squamous epithelium is replaced by a simple columnar lining which shows unusual intestinal characteristics (Peters et al., 2019). The cellular mechanism for how BO arises has remained unclear. Several theories have been proposed including transdifferentiation of the stratified squamous epithelium, re-population of the lower oesophagus *in situ* from an altered submucosal stem cell compartment (reviewed in Hayakawa et al., 2021), or more recently upward migration of stomach columnar epithelial cells from at or below the gastro-oesophageal junction (GOJ) (Polak et al., 2015; Nowicki-Osuch et al., 2021; Singh et al., 2021). In large part, the challenge of identifying the cell-of-origin stems from the cells in BO failing to replicate in totality any discrete aspect of the adult gastrointestinal tract. Recently, this led us to hypothesise that BO, and in turn OAC, might actually invoke reversion to a primitive state more reminiscent of embryonic and/or fetal development, from which variable components of misdirected gastrointestinal re-differentiation might arise. We discovered a gene regulatory network in BO and OAC centred on the critical developmental transcription factors (TFs), hepatocyte nuclear factor 4 alpha (HNF4A) and GATA-binding protein 6 (GATA6), neither of which are ordinarily expressed in the adult oesophagus (Rogerson et al., 2019). Indeed, HNF4A is capable of opening chromatin in squamous oesophageal epithelial cells to drive acquisition of a BO-like transcriptional signature (Rogerson et al., 2019), and associated phenotypic changes (Grimaldos Rodriguez et al., 2023). Others have since demonstrated that HNF4A is able to transform adult gastric epithelial cells into a BO-like state (Nowicki-Osuch et al., 2021; Singh et al., 2022).

Understanding whether BO might be a reversal of development has been blocked by rudimentary knowledge of oesophageal development in humans compared to other mammals. Structurally, the oesophagus first forms during the fourth week post-conception as the major part of the foregut endoderm between the buccopharyngeal membrane and the dilation that gives rise to the stomach (reviewed in Zhang et al., 2021). The TF, Sex determining region Y-box 2 (SOX2), is needed for oesophagus formation and marks the initial simple columnar epithelium (Que et al., 2007; Trisno et al., 2018). By inference from mouse, transition to the multi-layered epithelium is controlled at least in part by the TF TP63, from cells in the basal stem cell compartment of what eventually becomes stratified squamous epithelium (Rosekrans et al., 2015; Zhang et al., 2021). Further molecular details in humans are exceedingly limited. Therefore, in this study we set out to dissect the molecular development of the lower oesophagus at single cell resolution, and to impute the gene regulatory networks and trajectories responsible for the dynamic advance from simple columnar towards stratified squamous epithelium. The findings led us to undertake complementary studies to explore the relevance to the cellular mechanism underlying BO.

## RESULTS

### Morphology of human oesophageal development

We first wanted to define reliably which time points would allow us to capture the dynamic epithelial changes in the developing lower oesophagus. The epithelial layer converts from a simple columnar (‘primitive’) state, uniformly present at 7-8 weeks post-conception (wpc), to a ‘transitory’ epithelium captured at 9-12 wpc that now contains regions of several layers with a basal cell compartment and a luminal surface that appeared ciliated (Fig. 1A; Supplementary Fig. S1A). A more complex multi-layered epithelium comes to predominate (‘stratified’; noticeable by 15 wpc), still containing ciliated cells but which is known to become squamous over time and remains that way in the adult (Fig. 1A; Supplementary Fig. S1A and B). In contrast, the fetal stomach remained lined throughout by simple columnar epithelium. Consequently, a dynamic transition zone develops in the lower oesophagus leading into the stomach. Initially, this was broad, but narrowed during development. In the adult, it is sharply demarcated as the squamo-columnar junction and is most commonly coterminous with the GOJ (Supplementary Fig. S1B). Having defined when we could represent these different stages and features of human oesophageal development (primitive, transitory and stratified), we went on to dissect tissue from the lower oesophagus at 7, 9 and 15 wpc to study the individual cell types in molecular detail using a combination of single nuclear (sn) RNA-seq and ATAC-seq (Fig. 1B).

**Fig. 1:**
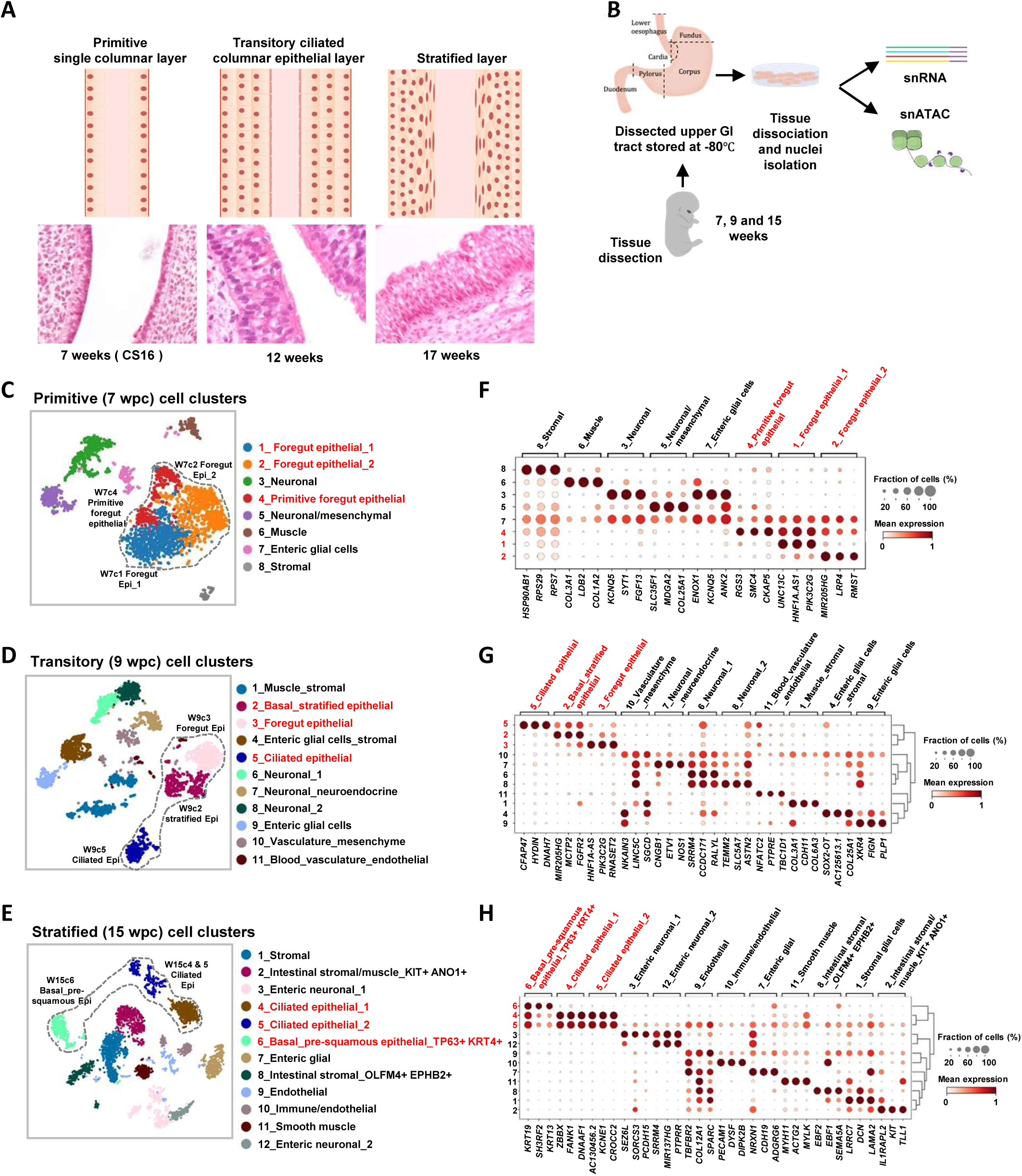
Single nuclear RNA-seq analysis of the developing human oesophagus. (A) Diagrammatic representations of the structures of the developing lower oesophagus. Representative H&E stained sections of each stage are shown below the diagrams. (B) The overall project workflow from tissue collection to sequencing. (C-E) tSNE plots of 7 (B), 9 (C), and 15 (D) week cell clusters based on snRNA-seq. Major epithelial populations are circled and annotated on the plots. (F-H) Dotplots of the relative average expression of three representative markers for each of the cell clusters found at weeks 7 (F), 9 (G) and 15 (H). The fraction of cells expressing each marker and relative expression levels (column normalised) are represented by the size and intensity respectively, of each dot.

### Cellular composition of the developing oesophagus

First, we performed snRNA-seq on 2,121, 2,972 and 4,615 nuclei at primitive (7 wpc), transitory (9 wpc) and stratified (15 wpc) stages respectively. At each time point, we visualised the different cell populations by UMAP and identified 8-12 distinct clusters of cells (Fig. 1C-E), which we annotated based on stage, overall gene expression profile and selected marker genes (Fig. 1F-H; Supplementary Fig. 1C-E; Supplementary table S1). For all three time points, some marker genes were predominantly found in a single cluster (e.g. *COL3A1* in cluster 6 and *SMC4*/*KIF23* in cluster 4 at 7 wpc; *EBF2* in cluster 8 at 15 wpc), while other genes distinguished a number of clusters (e.g. *GATA6.AS1* in clusters 1, 4 and 7 and *LRP4* in clusters 2 and 4 in primitive oesophagus at 7 wpc) (Fig. 1F-H; Supplementary Fig. 1C-E). In several cases, different clusters lacked unique marker genes and instead could only be deconvoluted by combinatorial gene expression. For example, at 15 weeks the two ciliated epithelial sub-populations were distinguished by *ZBBX*, *FANK1*, *DNAAF1*, *AC130456.2*, *KCNE1* and *CROCC2* and discriminated from each other by the presence (cluster 5) or absence (cluster 4) of a wide range of other genes common to other clusters (e.g. *COL12A1* and *SPARC*) (Fig. 1H).

To explore the evolution of cell types across developmental stages, we assembled all the data together into a single UMAP to reveal four broad cell lineages defined by different sets of marker genes: epithelial, mesenchymal, enteric nervous system and blood/vascular (Supplementary Fig. 2A-C). In keeping with the growing complexity of the oesophagus from its origin as a simple foregut tube, the proportion of epithelial cells declined with advancing fetal age (Supplementary Fig. 2D). By aggregating across stages, related cell populations could be visualised by the expression of single genes such as *PHOXA* in neuronal-related cells (Supplementary Fig. 2E). Similarly, different epithelial populations could be defined, such as *CFAP73*^+^ ciliated cells which are present at transitory and stratified stages but not in the primitive epithelium. Other epithelial populations showed overlapping, graded expression of marker genes; for example, combinations of *GATA6*^High^/*TP63*^Low^, *GATA6*^Intermediate^ ^(Int)^/*TP63*^Int^, and *GATA6*^Low^/*TP63*^High^ expression within and across clusters of developing epithelial cells (Supplementary Fig. 2A, B & E). *GATA6*^High^/*TP63*^Low^ cells were observed mostly in the primitive and transitory epithelium, but are near absent in the more complex stratified epithelia cells by 15 weeks. Conversely, *GATA6*^Low^/*TP63*^High^ cells characterised the basal cells of transitory and stratified epithelium at 9 and 15 weeks (Supplementary Fig. 2E). The changing composition of the epithelial cell populations as development proceeds, reflects the dynamic morphological changes taking place in the epithelium over this time period.

### Development of the oesophageal epithelial layer

To further understand the changes occurring during the development of the epithelial layer, the epithelial populations across the primitive, transitory and stratified stages were combined and re-clustered (Fig. 2A). Three major epithelial cell subtypes were identified, foregut columnar-like, ciliated and basal stratified cells (Fig. 2A, bottom). Hierarchical clustering or Pearson’s correlation consistently identified the *CFAP73*^High^ ciliated epithelial cells as the most distinct population (Fig. 2B-C and Supplementary Fig. 3A). Within the *CFAP73*^Low^ populations, the hierarchical ordering from the primitive columnar cells through to stratification was consistent with sequential *GATA6*^High^/*TP63*^Low^, *GATA6*^Int^/*TP63*^Int^ and *GATA6*^Low^/*TP63*^High^ expression (Fig. 2B-C and Supplementary Fig. 3A). Amongst primitive populations, RNA velocity analysis implied W4c4 cells were the most rudimentary cells with a clear flow to the other two primitive epithelial clusters (Fig. 2A, W7c4, and Fig. 2D; Supplementary Fig. 3B) but directional flow between clusters was difficult to discern at later timepoints. In contrast to the close clustering of all cells at the primitive stage, the cell clusters become well separated over time, reflecting the more defined cell types established as a stratified epithelial layer is created (Fig. 2A and 2D; Supplementary Fig. 3B). Indeed, a clear switch towards a stratified epithelium is apparent by 15 wpc with a basal layer established, and the luminal layer being lined with *FOXJ1*-positive ciliated epithelial cells (Supplementary Fig. 3C).

**Fig. 2:**
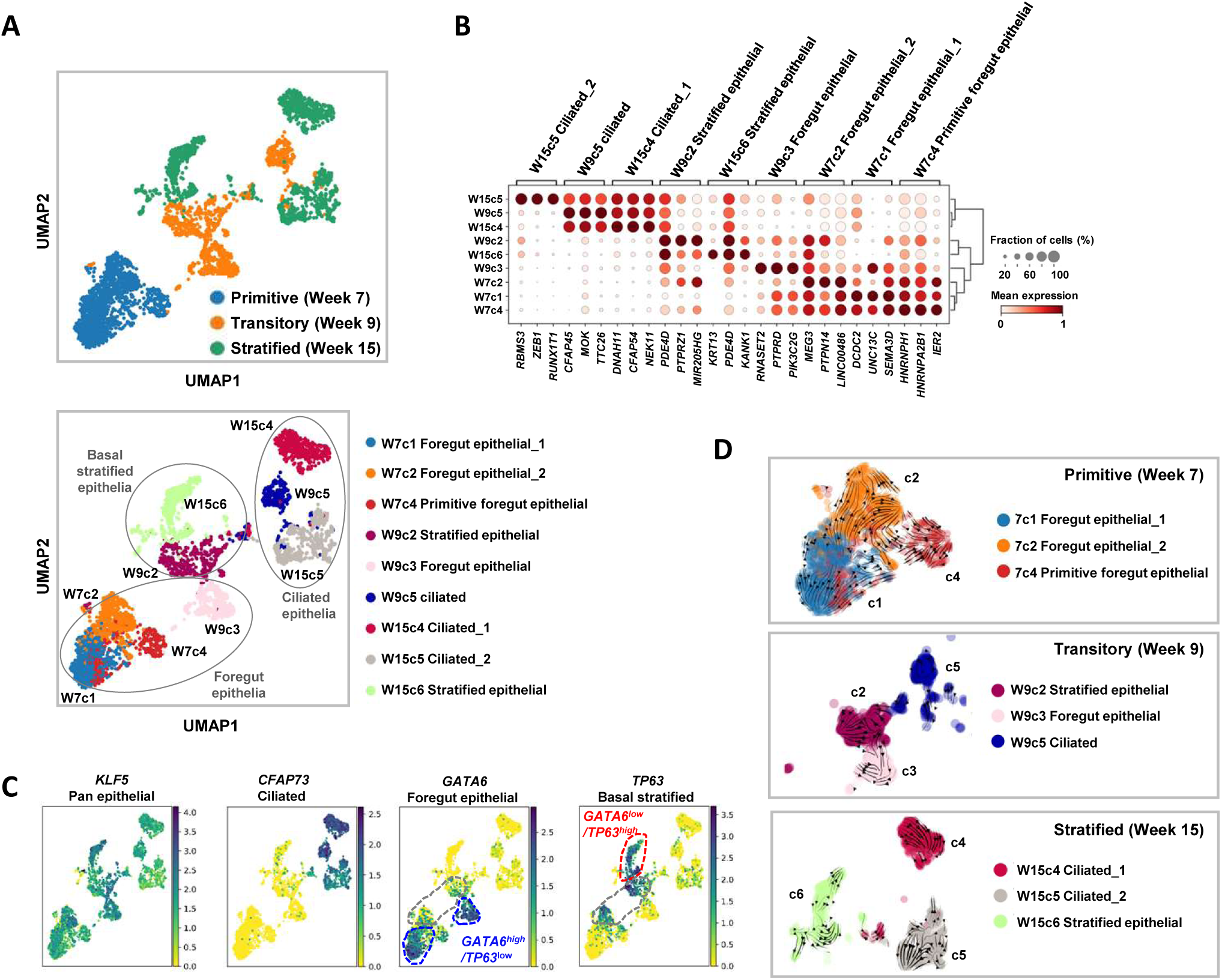
Development of the oesophageal epithelial layer. (A) UMAP of snRNA-seq data from the epithelial cell populations found during embryonic developmental stages at weeks 7, 9 and 15. Clusters are colour coded according to different epithelial cell types found at each developmental stage (top) or all epithelial cells at each time point (bottom). Major categories of epithelial cells are highlighted. (B) Dotplots of the relative average expression of three representative markers for each of the epithelial cell clusters found at different time points. The fraction of cells expressing each marker and relative expression levels (column normalised) are represented by the size and intensity respectively, of each dot. (C) Marker genes for each of the indicated epithelial cell subtype are shown superimposed on the UMAPs. GATA6^high^/TP63^low^ (blue), GATA6^low^/TP63^high^ (red) and intermediate GATA6^int^/TP63^int^ (grey) cell populations are outlined. (D) RNA velocity (scVelo) analysis of the epithelial cell clusters at each timepoint superimposed on tSNE plots.

Having begun to discern a likely differentiation pathway, we used a series of marker genes to check if any of the developmental epithelial populations are reminiscent of the corresponding adult anatomy, such as cells from the lower oesophagus, upper gastric epithelium as well as the BO metaplastic disease state (as defined in Nowicki-Osuch et al., 2021; Supplementary Fig. 3D). As expected, the basal stratified marker *TP63* was only detected in the normal oesophagus. In normal adult tissue, *GATA6* (foregut columnar maker) was expressed in some gastric epithelial populations, but these cells were unlike those found during development and in BO as they were largely *KLF5 and HNF4A* negative. *GATA6* was not detected in the normal lower oesophagus. *HNF4A* (another foregut columnar marker) was only expressed in the BO epithelium. All of the adult clusters lacked cilial markers, such as *CFAP73*, in keeping with the finding that ciliated cells are not generally detected in the adult oesophagus in North American populations (Scott et al., 2019). *GATA6*, *HNF4A* and *KLF5* collectively marked cells in BO and this pattern was not replicated amongst any of the adult populations examined but was reminiscent of the primitive and transitory epithelial sub-populations observed during human development.

Collectively, these data demonstrate that during development, the epithelial layer undergoes a dynamic transition from a largely simple primitive foregut in the embryo, through to three major cell types (primitive foregut, ciliated, and basal stratified epithelium) in the progression to the fetal stage, which subsequently resolves into the cells of the stratified epithelial layer, reminiscent of that found in the adult.

### Gene regulatory networks in the primitive and transitory epithelium

Given the growing evidence of a link between BO and oesophageal epithelial development, we decided to examine gene regulatory networks in our embryonic and fetal material to further understand the developmental pathways involved. We focussed on when the epithelial layer undergoes the most dynamic changes and performed snATAC-seq at the primitive (7 wpc) and transitory (9 wpc) stages on 4,612 and 481 nuclei respectively. At the primitive stage, twenty distinct clusters were identified (Supplementary Fig. 4A, left). All cell states previously identified in clusters in the snRNA-seq data (Fig. 1C) mapped by label transfer to the ATAC-seq-derived clusters except enteric glial cells (Supplementary Fig. 4A, right). Populations defined singularly by gene expression, for instance muscle cells, could be often be resolved as several distinct ATAC-seq clusters (in this case clusters 17, 18, 19 and 20; Supplementary Fig. 4A and B). This is suggestive of cells with similar gene expression profiles which may be in the process of reorganising their regulatory chromatin landscapes as they transition to different cell states. A similar phenomenon was observed for epithelial cells where four ATAC-seq clusters (1, 7, 8 and 10) mapped to the three populations defined by gene expression, allowing five distinct sub-populations to be discerned when the two modalities were integrated (Fig. 3A; right). These observations prompted us to re-cluster the epithelial populations in the RNA-seq data, which led to the identification of two minor and five major clusters with distinct sets of marker genes (Supplementary Fig. 4C-E). *HNF4*^+^ (cb) and *TP63*^+^ (cc) clusters could be discerned, along with an additional *MUC16*^+^ cluster (cd) and a cluster defined by the expression of mitotic cell cycle genes (ce) (Supplementary Fig. 4E). The latter is in keeping with our designation of these cells as epithelial progenitors, and is supported by RNA velocity analysis which shows a flow from these progenitors (ce) towards the *HNF4*^+^ (cb) and *TP63*^+^ (cc) clusters (Supplementary Fig. 4F).

**Fig. 3:**
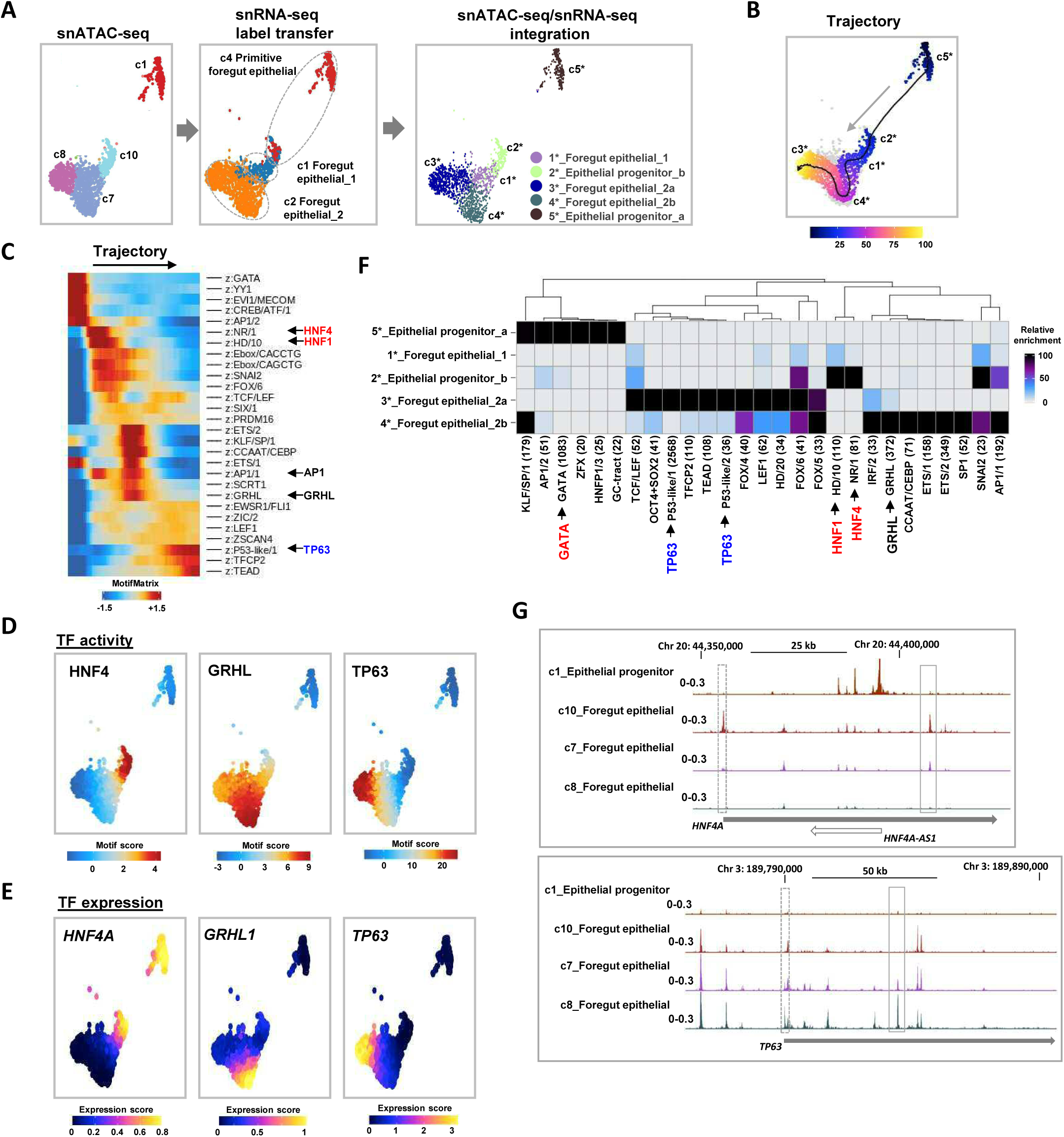
Development regulatory networks in primitive stage epithelial cell populations. (A) UMAPs derived from snATAC-seq data showing the epithelial cell populations in 7 week embryos. Cells are clustered according to snATAC-seq signal (left), annotated based on label transfer from snRNA-seq (middle) and reclustered based on combined use of snATAC- and snRNA-seq (right). (B) Trajectory analysis of week 7 epithelial cells from clusters c1-5* from the integrated analysis. (C) TF motif deviation score across the trajectory depicted in part (B). (D) Transcription factor binding motif scores for individual cells projected on the epithelial population ATAC-seq-derived UMAP. (E) Gene expression scores (from gene integration matrix) for the indicated transcription factors for individual cells projected on the epithelial population ATAC-seq-derived UMAP. (F) Heatmap showing the relative enrichment of the indicated transcription factor binding motifs in each of the epithelial cell clusters (scale bar shows enrichment scores in negative log_10_ of p_adj value). Motifs are given family names (Vierstra et al., 2020) and those discussed in the text are highlighted. (G) UCSC browser showing ATAC-seq peaks surrounding the indicated loci and original ATAC-seq-derived cell clusters. Promoter and intergenic/intragenic peaks changing between samples are highlighted by dashed and solid line rectangles respectively.

We next applied ArchR (Granja et al., 2021) to analyse DNA motif enrichments across a trajectory through the newly refined ATAC-seq clusters beginning with the putative progenitor cells in cluster c5* (Fig. 3B). The top variable motifs across the trajectory act as surrogates for TF activity along this trajectory (Fig. 3C). GATA motifs dominated the start of the trajectory, and cells then transitioned through states dominated by HNF4/HNF1, then AP1/GRHL and finally TEAD/TP63 TF activity. These findings are consistent with the differentiation path imputed purely from the snRNA-seq data with the transition from *TP63*^Low^ to *TP63*^Int^ and then *TP63*^High^ expression. These cell state transitions were clearly visible by projecting the motif deviation scores for HNF4, GRHL and TP63 family TFs onto the UMAP defined by ATAC-seq (Fig. 3D) and supported by the corresponding imputed snRNAseq gene expression data for predicted TF activity (Fig. 3E). The identification of *GRHL1* expression/GRHL activity demarcated a novel transitional cell population (c3*) between the *HNF4A*/HNF4^high^ and the *TP63*/TP63^high^ cell states. These findings were further substantiated by relative motif enrichments in each of the five newly defined ATAC-seq clusters, with HNF4, GRHL and TP63 motifs again figuring prominently in distinct clusters (Fig. 3F). We also applied SCENIC+ (González-Blas et al., 2023) to derive gene regulatory networks in the 7 wpc cell clusters and provided further support for the existence of foregut epithelial cell GRNs driven by members of the HNF4 and GRHL TF families, along with other co-associated GRNs driven by a range of TFs including GATA6, FOXA1 and KLF5 (Supplementary Fig. 5A and B). In keeping with the evolving chromatin landscapes, the changes in expression of prominent regulatory TFs, were mirrored by chromatin accessibility changes at their loci as exemplified by *HNF4A* and *TP63*. The *HNF4A*, promoter and putative intronic enhancer peaks from ATAC-seq cluster 10 were diminished in transitional cluster c7 and virtually absent in cluster c8 (Fig. 3G, top). Reciprocally, a promoter peak at the *TP63* locus (giving rise to the short ΔN-TP63 isoform) is absent in ATAC-seq cluster 10 but emerges in cluster c7 and is retained in c8, when it was accompanied by an additional putative intronic enhancer peak (Fig, 3G, bottom). Collectively, these data identify the key TFs that define different cell states in the primitive epithelium and underline HNF4A and GRHL1 as sequentially acting TFs on the way to dominant TP63 activity as cells progress from primitive *TP63*^Low^ towards *TP63*^High^ stratified cell states in the developing lower oesophagus.

We hypothesised that the switch from HNF4A to TP63 activity would be present and validated in transitory cells at 9 wpc as this stage is still prior to full stratification of the epithelium. Seven distinct clusters were identified at this stage by snATAC-seq, three of which were designated as epithelial by label transfer from snRNA-seq (Fig. 4A; Supplementary Fig. 6A). Combining the two modalities yielded four distinct epithelial sub-populations (Fig. 4A, right; Supplementary Fig. 6B). Ciliated cells remained as a distinct cluster but an additional intermediary group of cells (c2a) emerged with the snATAC-seq profile of primitive foregut epithelial cells and the snRNA-seq profile of basal stratified epithelial cells. A similar phenomenon was observed for other clusters, where combining modalities split several of the ATAC-seq clusters, typified by cluster 7, where the gene expression profiles identified three distinct subclusters, all related to neuronal cell states. This is suggestive of commonly derived cell types with similar epigenetic profiles which are in the process of transitioning to different cell states.

**Fig. 4:**
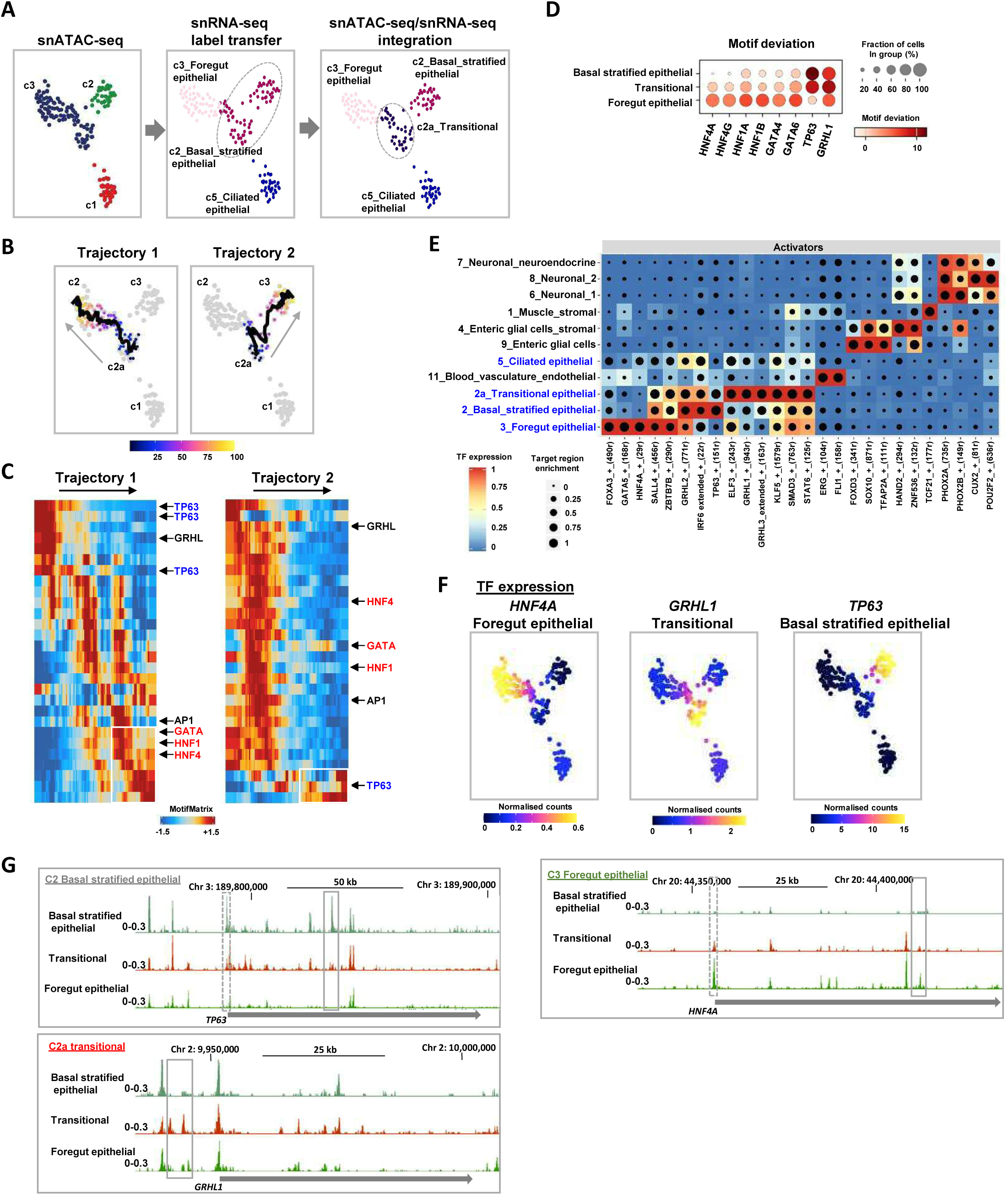
Developmental regulatory networks in the transitory epithelial cells. (A) UMAPs derived from snATAC-seq data showing the epithelial cell populations in 9 week embryos. Cells are clustered according to snATAC-seq signal (left), annotated based on label transfer from snRNA-seq (middle) and reclustered based on combined use of snATAC- and snRNA-seq (right). (B) Trajectory analysis of week 9 epithelial cells from clusters c2, c2a and c3. (C) Scaled TF motif deviation score across the trajectories depicted in part (B). (D) Dot plots showing the deviations in motif scores (indicated by colours) for the indicated DNA binding motifs in the indicated 9 week epithelial cell clusters (% of cells with open motif deviation score Indicated by size of dots). (E) Gene regulatory networks in 9 week cell clusters identified using SCENIC+. The top scoring transcription factor networks are shown (eRegulons correlation coefficient above 0.70 or below -0.65). (F) UMAPs showing the expression (gene integration scores from snRNA-seq) of the indicated genes. (G) UCSC browser showing ATAC-seq peaks surrounding the indicated loci and cell clusters. Promoter and intergenic/intragenic peaks changing between samples are highlighted by dashed and solid line rectangles respectively.

To identify potential developmental trajectories, we first re-clustered the snRNA-seq data associated with epithelial clusters c2 and c3 and applied RNA velocity analysis. Three major clusters were identified, with a newly identified transitional population which showed directional flow towards either foregut or basal stratified epithelial clusters (Supplementary Fig. S6C). Next, we used ArchR (Granja et al., 2021) to impart trajectories on the overlapping epithelial clusters, and based on RNA velocity analysis each started at the newly defined intermediary state (Fig. 4B). We examined the dominant TF motifs as surrogates for TF activity across these trajectories and found that cells at the end of trajectory 1 are characterised by high HNF4 and HNF1 activity whereas those at the end of trajectory 2 contain high TP63 activity (Fig. 4C; Supplementary Fig. 7A). In both cases, a transitional state was observed which commonly contained high GRHL activity (Fig. 4C; Supplementary Fig. 7A) as observed in the transitional state in the epithelial cell populations at 7 weeks (Fig. 3C-E). Plotting TF activity scores on top of the UMAPs (Supplementary Fig. 7B) or calculating TF activities using DNA binding motif deviation scores (Fig. 4D) reinforced the trajectory analysis. HNF4 activity was highest in c3 foregut epithelial cells, GRHL in c2a transitional cells, and TP63 in c2 basal stratified epithelial cells (Fig. 4D; Supplementary Fig. 7B). These findings were further strengthened by identifying cluster-specific TF motif enrichments for HNF4, GRHL, and TP63 and RFX in each of the four epithelial cell clusters (Supplementary Fig. 7C). The latter observation was confirmed by plotting motif deviation scores which localised RFX activity to the ciliated epithelial cell cluster in keeping with the known role for this transcription factor in specifying ciliated cell fates (Choksi et al., 2014). Moreover, GRN analysis identified regulons for a wider number of regulatory TFs for each cluster, and again uncovered the same categories of regulatory TFs with HNF4 in foregut, GRHL1 in transitional and TP63 in basal stratified epithelial cells (Fig. 4E). Moreover, TF co-regulatory activity was suggested from correlation analysis, which is particularly marked in the foregut epithelial cluster where HNF4A, GATA5 and FOXA3 regulons coincide (Supplementary Fig. S7F).

Further analysis showed that expression of members of each TF subfamily broadly follows their predicted activity profiles with high *HNF4A* expression in c3 and high *TP63* expression in c2 (Fig. 4F). *GRHL1* expression is high in the transitional cells located in the c2a cluster between c2 and c3. Similarly, *KLF5* shows highest expression in the transitional population, although this pan-epithelial TF is expressed in all epithelial cell clusters (Supplementary Fig. 7E). These changes in expression are mirrored by chromatin accessibility changes in the *TP63*, *GRHL1* and *HNF4A* loci where unique peaks emerge in each cluster and in the case of *HNF4A* and *TP63*, an opening and closing of the promoter region as cells progress from the transitional state (Fig. 4G).

In summary, these data demonstrate that during development, the epithelial layer undergoes a dynamic transition from a largely simple primitive foregut, through the production of three major cell types (primitive foregut, ciliated, and basal stratified epithelium), each defined by dominant regulatory TFs. Ultimately the simple columnar epithelial layer resolves into the stratified epithelial layer that typifies the adult oesophagus.

### Relationship between epithelial states in BO and early oesophageal development

Our observation that the primitive epithelial state of early development mimics features of the metaplastic transition of BO led us to deepen our search for shared molecular features. We interrogated published single cell (sc) RNA-seq data (Supplementary Fig. 9A; Nowicki-Osuch et al., 2021) and newly generated snATAC-seq data from a non-dysplastic BO sample (Supplementary Figs. 8A, B and C; Supplementary Fig. 9B). Fourteen cell clusters were identified from 6,762 nuclei in the new snATAC-seq data (Supplementary Fig. 9B). By label transfer from the scRNA-seq data, all of these clusters mapped neatly across with the exception of the small cluster, c10, which was therefore excluded from further consideration (Supplementary Fig. 9C and D). Amongst the epithelial populations (clusters c3-9 by snATACseq), label transfer pointed to differentiated populations of columnar, endocrine and goblet cells, as well as dynamically changing columnar populations, clearly demarcated by snATAC-seq but with shared or overlapping gene expression profiles (Supplementary Fig. 9B-D; clusters c5, 6 and 8). By UMAP, these three clusters were flanked left and right by undifferentiated (c7) and differentiated (c4) columnar populations, most likely implying connections through transitional states.

We recently demonstrated that a TF network containing members of the HNF4, HNF1, FOXA and GATA4/6 subfamilies broadly defined the core gene regulatory pathways in BO and that this network is retained in OAC (Fig. 5A; Rogerson et al., 2019; Chen et al., 2020). We hypothesised that the genes encoding these TFs might actually demarcate and be accountable for the different BO epithelial sub-populations. Indeed, their expression profiles overlap in these populations (Supplementary Fig. 9E) and together with activity, as imputed from DNA binding motif enrichment (Supplementary Fig. S9F), indicates a regulatory role for the GATA-HNF1-HNF4 axis in the transitional columnar undifferentiated dividing populations. These findings were further underlined by plotting imputed expression and motif deviation scores for TF activity on the snATAC-derived UMAPs where *GATA4*, *HNF1A* and *HNF4A* expression and their corresponding DNA binding motif deviation scores converge on the transitional undifferentiated epithelial BO populations (Supplementary Fig. 9G and H). High HNF4A activity is maintained in the differentiated BO epithelial cells. These findings reinforce the idea that the epithelium of BO is itself differentiating and that the GATA-HNF1-HNF4 TF axis plays a key role in this process. This obvious parallel to development led us to return to the epithelial subtypes of the developing oesophagus to see if we could detect evidence of expression and activity of members of the *HNF4*, *HNF1, FOXA* and *GATA4*/*6* subfamilies. The expression of each gene was predominant in the more primitive cell-types, but less apparent in differentiated basal or ciliated cells (Fig. 5B). Co-expression and overlapping imputed binding activity was highest in transitory foregut epithelial cells (cluster c3 at 9wpc; Fig. 5B-D; Supplementary Fig. 10A-B). Given these associations, we asked whether we could detect interconnections within the transitory foregut epithelial cell GRNs and found high connectivity among these TFs (Fig. 5E).

**Fig. 5:**
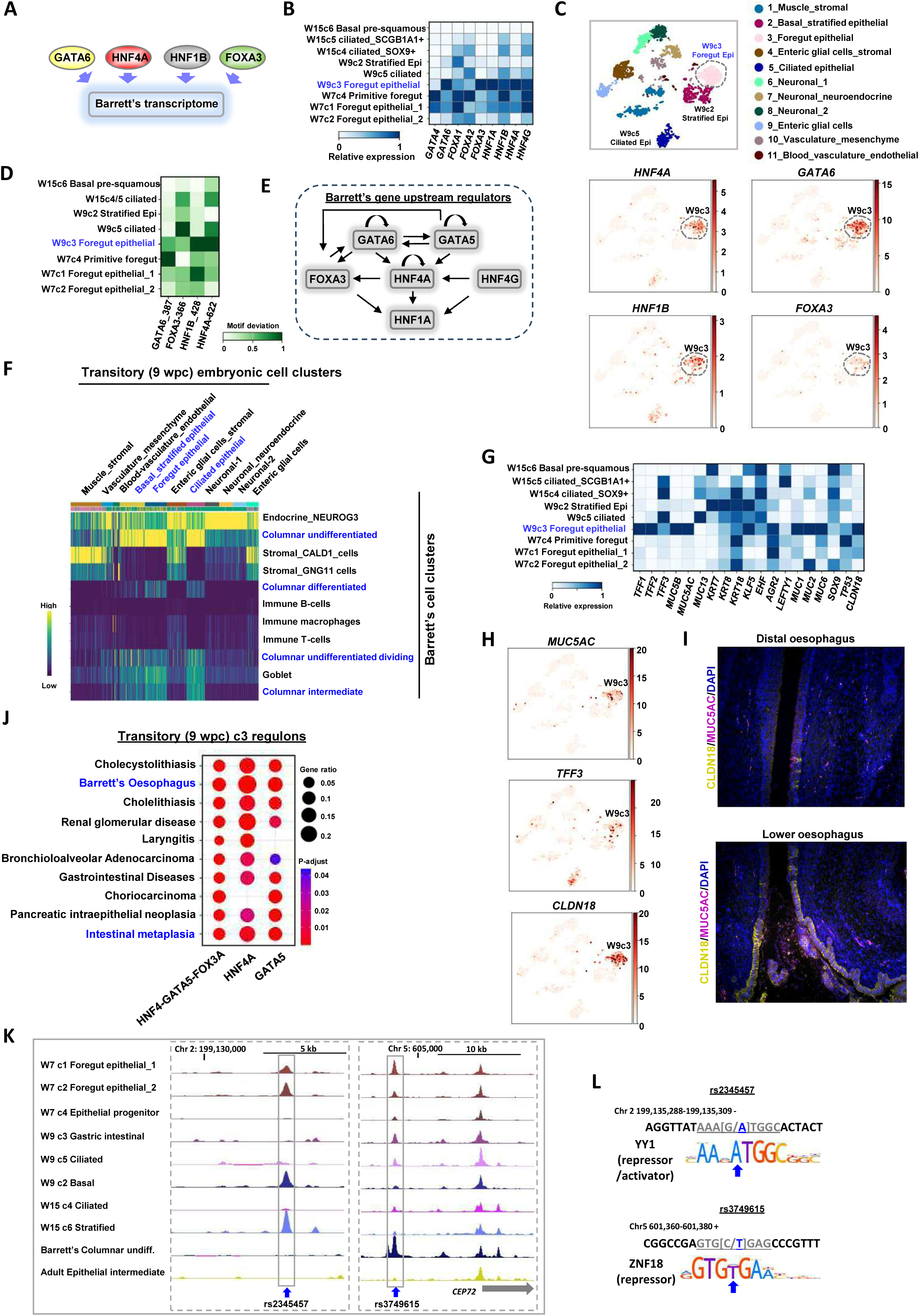
Barrett’s oesophagus resembles a transient developmental epithelial population. (A) Diagrammatic representation of the transcription factor repertoire driving Barrett’s cell identity (Rogerson et al., 2019). (B) Heatmap showing the relative expression (column normalised) of the indicated transcription factors in each of the developmental epithelial cell clusters from weeks 7, 9 and 15 (W7, W9 & W15). (C) tSNE plot showing all embryonic 9 week cell clusters (top) with the expression of the indicated transcription factors projected on top of these (bottom). (D) Transcription factor binding motif scores (column normalised) in open chromatin regions associated with each of the indicated epithelial cell clusters. (E) Regulatory links derived from the week 9 foregut epithelial cluster GRNs, depicting regulatory interactions between the core TFs. Arrows depict a directional regulatory link. (F) Heatmap showing similarity scores between each cell in the transitory stage 9 week developmental clusters (x-axis) and the corresponding cell types found in Barrett’s cell clusters (y-axis). (G) Heatmap showing the relative expression of the indicated Barrett’s marker genes in each of the embryonic epithelial cell clusters from weeks 7, 9 and 15 (W7, W9 & W15). (H) tSNE plot showing all embryonic 9 week cell clusters with the expression of the indicated Barrett’s associated marker genes projected on top of these. (I) IHC of MUC5AC and CLDN18 protein expression in week 9 embryos at the GOJ and distal part of the oesophagus. (J) Enriched DisGeNET GO terms for the genes in the combined HNF4A (29r), FOXA3 (490r) and GATA5 (168r), and individual HNF4 and GATA5 regulons in 9 week c2 foregut epithelial cells. (K) UCSC genome browser view of ATAC-seq signals surrounding SNPs rs2345457 (left) and rs3749615 (right) in the indicated clusters of developmental epithelial cells, Barrett’s undifferentiated columnar epithelia and adult oesophageal epithelial cells. The arrow below the right track represents the directionality and extent of the *CEP72* gene covered by the views. Peaks containing the SNPs are boxed. (L) Sequences around the significantly associated GWAS SNPs rs2345457 (top) and rs3749615 (bottom). The genomic location and DNA strand is shown above the sequence. Base changes are shown as WT/risk allele and the risk allele is coloured blue. Logos for the DNA binding motif of YY1 and ZNF18 are shown and the bases in the corresponding DNA sequence are underlined. Arrow indicates the base changed by the risk SNP.

Based on these clues from key TFs, we looked more globally at potential similarities in expression between BO cell populations and the developmental epithelial clusters. Undifferentiated columnar cells in BO showed strong similarity to epithelial populations at primitive (Supplementary Fig. 10C) and transitory (Fig. 5F) stages. More specifically, well-recognised markers of BO, for both undifferentiated and differentiated columnar cells (Supplementary Fig. 9I), were expressed in greatest number in the transitory foregut epithelial cells at 9wpc (Fig. 5G, cluster c3). This strong overlap with cluster c3 is exemplified by *MUC5AC*, *TFF3* and *CLDN18* (Fig. 5H), with TFF3, MUC5AC and CLDN18 proteins detected in the lower oesophagus adjacent to the GOJ (Fig. 5I; Supplementary Fig. 10D). Furthermore, highly connected GRNs centred on the core BO TF network were observed in the transitory foregut epithelial cells (Supplementary Fig 11A), including links to many well-known markers of BO (Supplementary Fig. 11B). More generally the disease gene ontology (GO) terms “Barrett’s oesophagus” and “intestinal metaplasia” figured prominently within individual and combined regulons for the HNF4A, FOXA3 and GATA5 transcription factors (Fig. 5J; Supplementary Fig. 11C-E).

Together these data identify epithelial cell types in the oesophagus of the developing embryo and fetus which have very similar dominant TF regulatory networks and gene expression profiles to those in BO epithelial cells. These developmental cell types are prevalent transiently as the oesophagus transitions from a columnar to a stratified epithelium.

### Relationship of epithelial cells in BO to the developing stomach

Comparison of BO to the developing stomach alongside distal oesophagus is important, given growing evidence implicating gastric epithelial cells as a cell of origin of BO (Polak et al., 2015; Singh et al., 2021; Nowicki-Osuch et al., 2021). Therefore, we checked whether the developing stomach at a comparable stage to when we found the BO-like transitory oesophageal foregut columnar cells might also contain cells that resembled BO. We did this by combined snRNA- and snATAC-seq analysis (‘multiome’) for paired lower oesophagus and proximal stomach samples. The oesophageal data validated the previous analysis at 9 wpc, by identifying the same transitory epithelial populations, including foregut-like epithelial cells (clusters 2 and 3), alongside ciliated and basal cells (Supplementary Fig. 12A-D: Supplementary Table S2). By motif deviation scores, the same regulatory pathways driven by HNF4, GATA and HNF1 (foregut epithelial, cluster c3) and TP63 (basal, cluster c1) were also imputed (Supplementary Fig. 12E and F).

Parallel analysis of the stomach identified 12 clusters of cells (Supplementary Fig. 13A), each with a defining set of marker genes (Supplementary Fig. 13B-C; Supplementary Table S2), which for epithelial populations included foveolar (*FUT9*^+^; cluster 2), parietal (*CBLIF*^+^; cluster 4) and enteroendocrine (*ST18*^+^; cluster 1). Parietal/chief cells (cluster 3) shared overlapping markers, such as *FGD4*, with cluster 4 (parietal cells) suggesting an intermediary state. The clusters were also distinguished by strong enrichment for a range of imputed TF binding matched by gene expression, exemplified by ELF3/*ELF3* in foveolar cells (cluster 2), ESRRG/*ESRRG* in parietal cells (cluster 4) and RFX6/*RFX6* in enteroendocrine cells (Supplementary Fig. 13D-F).

We then compared the epithelial populations from the stomach and oesophagus. Oesophageal and gastric cell clusters segregated discretely by anatomical location, apart from some similarity between the gastric parietal cell populations (clusters 3 and 4) and the oesophageal foregut epithelial cells (cluster 2) (Supplementary Fig. 13G). However, when it came to similarities with BO, clustering was with oesophageal primitive and transitory foregut-like epithelial cells; in particular, the more primitive population (7 wpc) with undifferentiated BO columnar cells and transitory cells (9wpc/multiome) and the more differentiated BO columnar cells (Supplementary Fig. 14A). Conversely, there was little similarity between BO and any of the gastric populations (or the more differentiated oesophageal cell types). Narrowing in on our BO marker genes (Supplementary Fig. 9I) revealed very similar results for the transitory foregut-like oesophageal cells (cluster 3), with high differential expression for many of the key genes (Supplementary Fig. 14B). In contrast in the stomach, only the foveolar cells (cluster 2) showed lower level expression for a subset of Barrett’s marker genes (Supplementary Fig. 14B and C). Genes encoding TFs for the core BO regulatory network were also co-expressed in oesophageal foregut-like epithelial cell clusters, but tended to be distributed individually across different gastric cell populations, implying an inability to function as a TF network (Supplementary Fig. 14D-E). Taken together, these data point to gene expression in the developing stomach being poorly aligned to that of BO compared to very strong parallels globally, within the BO marker gene set or known TF networks, with the transitory oesophageal foregut-like epithelial populations.

### Barrett’s susceptibility loci map to developmental enhancers in the oesophageal epithelial cell populations

Genome-wide association studies (GWAS) have identified 27 sequence variants associated with increased risk of BO and/or OAC, many of which reside in non-coding regions (Schroder et al., 2022). Most of these have not been followed up functionally to identify the causal gene, in part because of a lack of access to RNA-seq and ATAC-seq data from relevant tissue types. It is plausible some of the variants impact susceptibility to loss of the normal stratified squamous phenotype; however, others would be predicted to drive acquisition and maintenance of the BO epithelium. Having shown that BO adopts a developmental oesophageal phenotype (rather than metaplasia from one adult cell-type to another), we predicted that some BO GWAS non-coding variants would reside in imputed regulatory elements in the epithelial cell populations during oesophageal development. Therefore, we overlaid the embryonic/fetal-specific accessible chromatin peaks with the locations of SNPs reaching suggestive significance in the BO/OAC GWAS (P-values <1×10^-5^) and excluded peaks that are also found in adult epithelial cells. Ten of these SNPs (of which 4 reach genome-wide significance (<5 x 10^-8^)) overlapped with 8 open chromatin peaks in the developmental epithelial cell clusters (Supplementary Table S3). These 10 SNPs mark 8 loci associated with BO/OAC (2 SNPs are in strong LD (r2>0.9) with one other SNP within the set of 10). Several are in strong LD with previously identified lead SNPs from the BO/OAC GWAS and are associated with regions encoding *SATB2*, *CEP72*, *CFTR* and *MFHAS1* (Supplementary Table S3; Chen et al., 2022; Schroder et al., 2022). We looked at several in detail; all aligned with open chromatin peaks from different combinations of the developmental clusters, with five of them present in the primitive foregut-like epithelial clusters (Fig. 5K; Supplementary Fig. S15A). Two of these SNPs are located in peaks unique to developmental epithelial cell clusters (rs2345457 and rs55982826) whereas the other four SNPs are located in peaks also found in the undifferentiated columnar epithelial cells in BO (Fig. 5K; Supplementary Fig. 15A). Moreover, the SNPs are all predicted to result in reduced activity of the imputed regulatory element, either through the creation of potential repressor binding sites (rs2345457, rs3749615 and rs1264706; creating YY1, ZNF18 and ZNF708 binding elements) or loss of binding of a transcriptional activator (rs55982826, rs17451754 and rs907183; disrupting KLF5, NFIC and TEAD binding elements, respectively)(Fig. 5L; Supplementary Fig. 15B).

In summary our results have uncovered profound similarities in gene expression and in TF regulatory networks between the columnar epithelium of BO and a transient population of oesophageal but not gastric epithelial cells during development; and that genetic risk variants for BO likely influence the activity of regulatory elements which are transiently active in these epithelial cell populations. In combination, these data support the theory that BO, as a pre-malignant state to OAC, arises directly from reversion of cells in the adult upper GI tract to a transient state found in epithelial cells of the developing human oesophagus.

## DISCUSSION

Towards the end of the first trimester of human development, the developing oesophagus undergoes an intricate conversion from a columnar to a squamous epithelial lining. In its lower portion this event is intertwined with the creation of the squamocolumnar and gastro-oesophageal junction with the stomach. Here, we have used a combined approach of profiling gene expression and chromatin accessibility at the single cell level to define the cell types that are present in the lower oesophagus and the underlying dominant TFs that specify their phenotypes. Harnessing this information, we discovered that in the disease state, BO, the metaplastic columnar epithelial cells show similarities in their gene expression profiles and regulatory TF networks to a transient epithelial population present in early development before and during the transition to a squamous epithelium. Strikingly, BO is generated by opposite transition from a squamous to columnar epithelium suggesting that normal development and the mechanism underlying BO might be highly similar processes operating in reverse directions. Our data therefore provide a molecular rationale for this dynamic transition whereby reawakening of the dominant regulatory networks found during development results in re-establishment of an embryonic-like columnar epithelial layer. By extension, this implies a residual epigenetic memory of the developmental state in adult cells in the upper GI tract that is susceptible to reactivation following injury to the oesophageal lining. Indeed, such reversion to a fetal-like state appears to be a common event across the intestine as a regenerative response to damage and inflammation (Viragova et al., 2024).

The diversity of cell types in BO are not all developmental in appearance. During development, the oesophageal layer transitions from a series of primitive columnar-like cells in the embryo through to an initial mixed population of ciliated, columnar and squamous-like cells. As development progresses, the columnar-like cells, and eventually the ciliated cells, are progressively lost to leave the adult stratified squamous epithelium. The columnar-like cells at primitive stages exhibit imputed activity of TFs from the HNF4, HNF1, GATA and FOXA families. This same network of TFs is re-activated in BO epithelial cells (Rogerson et al., 2019). Commonalities are apparent as HNF4A expression and imputed activity increases as cells become more specified in both development and BO. However, the relative expression of different members of these TFs varies between development and BO. Classical BO marker genes, such as *MUC5AC*, *TFF3* and *CLDN18*, were also unevenly expressed amongst the different epithelial populations both in BO and in development. This lack of complete congruence is unsurprising as any reversion to a developmental state would most likely be imprecise and the presence of other cell-types, such as endocrine cells, could well indicate additional aspects of re-differentiation that occur in BO. Moreover, as both the gastric and oesophageal epithelium arise from foregut endoderm, there may be yet as uncharacterised endodermal cell types from earlier developmental timepoints that may more closely resemble BO. Indeed, as BO and gastric cardia metaplasia are being increasingly considered as molecularly similar entities (Nowicki-Osuch et al., 2023), such a scenario appears entirely possible.

During development, TP63 expression and imputed chromatin binding activity were detected at the primitive stage in advance of a fully stratified squamous epithelium, suggesting that the TF’s regulatory function is initiated at a relatively early developmental timepoint. Only later does TP63 manifest its full activity to drive the gene expression programmes characteristic of the squamous epithelial state. Our detection of RFX TF activity in ciliated epithelial cells marked by *FOXJ1* expression is consistent with both playing known roles in specifying ciliated epithelial cell states (Choksi et al., 2014). In addition to the defined epithelial populations, we also uncovered transitional populations, often expressing mixed columnar and squamous cell markers. It is not clear how these transitions are controlled but at both the early primitive and transitory stages, the transitional populations exhibit high GRHL TF family activity consistent with previous studies that demonstrated a role for these TFs in priming epithelial enhancers for subsequent activation by lineage/cell type-specific TFs (Jacobs et al., 2018).

Several different mechanisms have been proposed for the development of BO based on different cells of origin located in the adult gastric cardia, GOJ and oesophagus (Leedham et al., 2008 Owen et al., 2018, Hu et al., 2007, Quante et al., 2012, Wang et al., 2011, Jiang et al., 2017, Hutchison et al., 2011, Nowicki-Osuch et al., 2021). While our work does not refute or confirm any of these locations as the definitive source of BO cells, the high similarity of BO populations to early oesophageal columnar epithelial cells (but not gastric) suggests the epigenetic memory of these developmental cells is maintained in whatever adult cell-type undergoes reversion in BO. Moreover, we also find that four out of 27 GWAS significant SNPs associated with susceptibility to BO/OAC map to open chromatin regions representing potential cis-regulatory elements in oesophageal epithelial populations during development. This suggests that susceptibility to BO could have its roots in imperfect assembly and/or function of the oesophageal epithelial layer during development.

In summary, our work provides the first detailed cell atlas of the developing human oesophagus and provides novel insights into the regulatory networks specifying the formation and transition of the epithelial lining. We uncovered evidence for BO-like cell populations during embryonic epithelial development with important implications for understanding how BO is specified. Ultimately, as BO is a pre-malignant state, extending this knowledge derived from human development in utero may unlock fresh mechanistic understanding for malignant transformation to OAC, which is currently lacking.

## MATERIALS and METHODS

### Sample dissection

Human embryonic and fetal material was collected under ethical approval from the North West Research Ethics Committee (23/NW/003), informed consent from all participants and according to the Codes of Practice of the Human Tissue Authority. Tissue collection took place on our co-located clinical academic campus overseen by our research team ensuring immediate transfer to the laboratory. Embryonic material was staged by the Carnegie classification (O’Rahilly and Muller, 2010). Fetal material (after 56 days post conception) was staged by foot length and ultrasound assessment. Individual tissues and organs were immediately dissected in cold phosphate buffered saline. In brief, the lower third of the oesophagus was dissected to the gastro-oesophageal junction, and the upper fundus was dissected from the stomach. All visible adherent mesenchyme was removed under a dissecting microscope. Non-dysplastic BO samples were obtained from biopsies during routine endoscopy. In all cases, tissue was flash frozen using dry ice / isopropanol.

### Immunohistochemistry

Immunohistochemistry and haematoxylin and eosin (H&E) staining was performed in both developmental and adult tissue as described previously (Jennings et al 2013, Jennings et al 2017), and using the primary antibodies listed in Supplementary Table S4. Briefly, human embryos and fetuses were fixed within 1 hr in 4% paraformaldehyde (PFA), processed, and embedded in paraffin wax for orientated sectioning at 5 μm intervals before downstream H&E and immunohistochemical staining.

### Single nucleus RNA-seq, ATAC-seq and multiome (ATAC and RNA) library construction

For single nucleus RNA- and ATAC-sequencing, nuclei were isolated from frozen tissue using the demonstrated 10X Genomics protocol (CG000124). Briefly, tissue was thawed before further dissection. Dissected tissue was incubated in chilled lysis buffer (10 mM Tris HCl [pH 7.4], 3 mM MgCl_2,_ 10 mM NaCl, 0.1% Tween 20, 0.1 % NP40 substitute, 0.01 % digitonin and RNAse inhibitor [0.2 u/μl]), on ice with gentle pipette and vortex mixing. Incubation time was adjusted according to developmental /adult stage. Cells were strained using 40 and 20 μm filtration, before centrifugation at 500x g for 10 minutes and further washing in wash buffer (10 mM Tris HCL [pH 7.4], 3 mM MgCl_2,_ 10 mM NaCl, 0.1 % Tween 20, 0.1 % BSA and RNAse inhibitor [0.2 u/μl]). Nuclei were resuspended for downstream RNA and ATAC library preparation using PBS, 0.1% BSA, RNAse inhibitor and nuclei buffer (10X Genomics Inc. Pleasanton, USA) respectively. For multiome analysis, nuclei were isolated from frozen tissue using the demonstrated 10X Genomics protocol (CG000375), using reagents outlined above. Nuclei were assessed for quality and quantity using a Countess II automated cell counter. Libraries were constructed using the following kits (10X Genomics): Chromium Next GEM Single Cell 3’ Reagent Kits v3 (RNA-Seq); Chromium Next GEM Single Cell ATAC Reagent kit v1.1 (ATAC-Seq); Chromium Next GEM Single Cell Multiome ATAC+ Gene Expression kit (multiome analysis). Libraries were processed on the Illumina NextSeq 500 platform.

### snRNA-seq analysis: Read mapping to genome

The sequence files generated from the sequencer were processed using the 10x Genomics custom pipeline Cell Ranger v3.0.1. This pipeline generated fastq files which are then aligned to the hg38 custom genome with all the default parameters of cellranger. The pipeline then identified the barcodes associated with cells and counted UMIs mapped to each cell. Cell Ranger uses STAR aligner to align reads to the genome so it discards all the counts mapping to multiple loci during counting. The uniquely mapped UMI counts are reported in a gene by cell count matrix represented as sparse matrix format. We aggregated all QC filtered cells in the different embryonic time-points to generate the aggregated dataset.

### snRNA-seq analysis: Cell quality filtering

Low-quality cells were removed from the dataset to ensure that the technical noise does not affect the downstream analysis. We used three parameters for cell quality evaluation, the number of UMIs per cell barcode (library size), the number of genes per cell barcode and the proportion of UMIs that are mapped to mitochondrial genes. Cells that had lower UMI counts than one Median Absolute Deviation (MAD) for the first two metrics and cells with higher proportion of reads mapped to mitochondrial genes with a cutoff of two MADs were filtered out. We then created violin plots for these three metrics to see whether there are cells that have outlier distributions which can indicate doublets or multiplets of cells. We removed outlier cells that have total read counts more than 50,000 as potential doublets. After these filtering steps, 9,708 cells (2,121 cells from Week 7; 2,972 cells from Week 9; 4,615 cells from Week 15) remained for downstream analysis.

### snRNA-seq analysis: Gene filtering and normalisation

Genes with average UMI counts per cell below 0.01 were filtered out when working with individual time-points as we assume these low-abundance genes do not give much information and are unreliable for downstream statistical analysis (Bourgon et al., 2010). In order to account for the various sequencing depth of each cell, we normalised the raw counts using the deconvolution-based method (Lun et al., 2016). In this method, counts from many cells are pooled together to circumvent the issue of higher number of zeros that are common in single cell RNA-seq data. Each cell is then divided by a pool-based size factor that is deconvoluted from the pool of cells and then multiplied by a million. These normalised data are then log-transformed with a pseudo-count of 1 added to all counts.

### snRNA-seq analysis: Visualisation and clustering

The first step for visualisation and clustering is to identify the Highly Variable Genes (HVGs). We first decomposed the variance of each gene expression values into technical and biological components using *modelGeneVar* function from *scran* and identified the genes for which biological components were >0.5 and with FDR value <0.05. We call these genes HVGs. These HVGs were then used to reduce the dimensions of the dataset using PCA. The dimensions of dataset were further reduced to 2D using t-SNE or UMAP, where the first 14 components of the PCA were given as input.

The cells at each developmental timepoint were grouped into their putative clusters using the dynamic tree cut method (Langfelder et al., 2008). Instead of choosing a fixed cut-point for the tree, the dynamic tree cut method identifies the branch cutting point of a dendrogram based on the underlying data and combines the advantage of both hierarchical and K-medoid clustering approach.

We generated an aggregated dataset by combining all cells from the three time points. Then, for this aggregated dataset we used SCANPY (Wolf et al., 2018) to generate UMAPs for visualisation. For this we first generated the HVGs with min_mean=0.0125, max_mean=3, min_disp=0.5. These HVGs were used to calculate the principal components (PC) of the data. We took 40 of the PCs and generated a neighborhood graph using the pp.neighbors() function. These neighbors were used to generate the PAGA graphs which are then used to calculate the UMAPs using the sc.tl.umap() function.

### snRNA-seq analysis: Identification of marker genes

To identify the marker genes for a cluster, we compared that cluster with all other clusters. We then report the genes that are differentially expressed in that cluster as the marker for the cluster. We used *rank_gene_groups* function from SCANPY package and conducted wilcoxon test to find these marker genes. These marker genes were then used to annotate the cell types of a cluster.

### RNA-velocity

We applied scVelo (Bergen et al. 2020) to identify transient cellular dynamics in each of our developmental stages. To calculate the spliced vs un-spliced ratio for each gene we used RNA-velocity’s command line tool, velocyto 10x with default parameters and used scVelo to compute velocity based on the spliced vs un-spliced ratios.

### snATAC-seq Analysis

We used cellranger-atac 2.0.0 to map fastq files to the hg38 genome. In summary, cellranger-atac trims primer sequences from the reads and maps these trimmed reads to the genome using modified version of BWA-mem (Li, 2013). After removing the PCR duplicates it then uses a similar algorithm to ZINBA (Rashid et al., 2011) to call peaks. Cellranger-atac then uses reads mapping to these peaks to identify signal from noise and uses it to call cells from background.

For our downstream analysis we used ArchR tool (Granja et al., 2021). After filtering out the doublets we did cell filtering based on the TSS enrichment score and minimum fragments in a cell. Cells having a TSS enrichment score of 4 and a minimum of 1000 fragments passed through our filter. After the QC we had 6,762 cells from Barrett’s, 1,658 cells from the normal adult oesophagus, 781 cells from week 15, 4,688 cells from Week 7 and 481 cells from week 9 embryonic/fetal oesophagus. We used Latent Semantic Indexing (LSI) for dimensionality reduction and used graph-based clustering to cluster the cells. Labels from cell type annotation in snRNA-seq were transferred to snATAC-seq to annotate the clusters with addGeneIntegrationMatrix that uses Seurat’s CCA method to do the label transfer. Peaks on these cell-types were then called using MACS2 (Liu et al., 2014) using the parameter *--call-summits --keep-dup all -- nomodel --nolambda --shift -75 --extsize 150 -q 0.1*. After identifying the marker peaks for each cell-type, they were then used to do motif enrichment. First, we used getMarkerFeatures function from ArchR to identify marker peaks. This function compares each peak in a cell group against its own background group of cells to determine whether the peaks have significantly higher accessibility. To create this background for a group of cells, ArchR first identifies the nearest neighbor cells in the multidimensional space that are not part of this group. Peaks that are enriched in a cluster with log-fold change more than 1 against its background group of cells with FDR<=0.01 are annotated as marker peaks for the cluster. *peakAnnoEnrichment* function was then used to analyse these groups of marker peaks for enriched binding sites of transcription factors. We use Cis-BP, a catalogue of direct and inferred sequence binding preferences as the motifset database. TF binding motifs were grouped according to similarity and annotated based on their (sub)families for concisely annotating figures (Vierstra et al., 2020).

### GWAS analysis with epithelial clusters

We first sub-setted all the epithelial cells from our all aggregated dataset. We then used MACS2 (Liu et al., 2014) to call peaks from the snATAC-seq data on these epithelial cells. This gives a single peak set for all the epithelial cells. We used this peak set for all our differential accessibility (DA) tests in this section. For DA, we first grouped the cells into two groups, all the developmental and BO epithelial cells constitute group-1 and all the normal adult epithelial cells constitute group-2. We then identified the peaks that are differentially accessible in group-1 compared to group-2. ArchR calls peaks using MACS2 and then uses an iterative merging strategy to merge the overlapping peaks into a single larger peak. ArchR also uses fixed-width of 501 bp for each peak. We used the getMarkerFeatures function from ArchR to calculate the differential accessibility. As mentioned earlier, ArchR uses a bias-matched group of background cells to compare and calculate the differential Accessibility. For statistical significance we used the Wilcoxon test which tests whether the distribution of peak opening across two groups are significantly different. We first used a threshold of mean difference more than 0.03 between group-1 and group-2 or FDR less than 0.01 to identify the initial peaks. This mean difference is calculated as the differences of the mean of peak accessibility distribution for a peak in two groups. Peaks passing this threshold were then further filtered to keep the peaks that have log fold change of more than 2.75 in group-1 compared to group-2. Peaks passing these thresholds were then overlapped with the GWAS SNPs and we removed the peaks that do not overlap with a SNP. For selecting the SNPs, we only choose those that have p-value less than 1×10^-5^ in the most recent OAC/BO GWAS meta-analysis summary statistics (n=5,563; Schroder et al., 2022). We lifted over the SNP positions from build 37 to build 38 for comparison with our snATAC-seq data. We consider a SNP to be overlapping if it is located within group-1 peaks and differentially accessible compared to normal adult epithelial cells. These overlapping peaks are then checked against their peak score which is calculated by MACS2 during peak calling as -log_10_(qvalue) where q values show the significance of the peak. Any peak that has a score lower than 25 are filtered out. Finally, we filtered out any peaks that have a score lower than 10 in the developmental cells only. This makes sure we do not have peaks that are only high in Barett’s without also being detected confidently in developmental cells. For this last filtering step we used peaks that were only called on the developmental cells only.

### Multiome data analysis

We used cellranger-arc 2.0.2 to pre-process our single cell multiome (ATAC and gene expression) sequencing data which involves performing alignment of the fastq files to hg38 genome, filtering, counting the barcodes, peak calling and counting of both gene expression (GEX) and ATAC modalities. For downstream analysis, we again used ArchR (Granja et al., 2021) as our core multiome analysis framework. We assessed the quality of the cells based on both modalities and filtered out the cells that have TSS Enrichment ratios lower than 4 or have mapped ATAC-seq fragments lower than 1,000 reads. We also used the ATAC modality to identify potential doublet cells. To infer these doublet cells, ArchR first synthesises in silico doublets by mixing reads from thousands of different combinations of cells. Cells matching the profile of these synthetic doublets are marked as potential doublet cells which are then filtered out. We then used the GEX modality to additionally filter out cells that have <500 reads mapped to them or cells that have higher than 5% of mitochondrial reads. After these filtering steps we retained 2,633 cells in gastric and 1,240 cells in oesophagus multiomes. We then applied iterative LSI both for individual modalities as well as for joint modalities where ATAC and GEX were joined using a weighted nearest neighbour algorithm (Hao et al, 2021). UMAP (Becht et al, 2018) plots were then generated by applying the algorithm individually on the LSI of ATAC modality, the GEX modality and the combined modality (GEX and ATAC). We then used graph clustering on these three LSI reduced dimensions to get clusters for the snRNA-seq modality, the snATAC-seq modality and for the joint modality. In addition to the gene expression that we obtained from GEX, we also calculated the gene score for the ATAC modality which is calculated by looking at the mapped reads located 100 kb either side of the gene start. Both the gene expression and gene score were used to annotate the clusters. We use MACS2 (Zhang et al., 2008) to call peaks for each of the ATAC-derived clusters. For our analysis we used the joint modality clusters for both the datasets. We use a fixed width of 501 bp for the peaks to facilitate downstream computational analysis without needing to normalise for peak length. We ran the Wilcoxon test to identify marker peaks for each cluster from this peakset and performed motif enrichment analysis as described above for snATAC-seq data, on these marker peaks. We also calculated the motif deviation for each of the motifs that explains how the accessibility of a motif deviates in a cell from its average accessibility across all cells. We then conducted TF footprinting to predict the precise binding location of a TF using ArchR (Granja et al., 2021). This combines the Tn5 insertions across many instances of a predicted TF binding site to address the requirements of higher sequencing depth for identifying a footprint.

We also inferred developmental trajectories using ArchR (Granja et al., 2021) which orders cells across a lower N-dimensional subspace based on a user-defined trajectory backbone.

### Inferring gene regulatory networks (GRNs) using SCENIC+

We used SCENIC+ (version 1.0.1.dev4+ge4bdd9f; González-Blas et al., 2023) to infer GRNs using both snRNA-seq and snATAC-seq. SCENIC+ links candidate enhancer regions with binding motifs to their candidate target genes. As input, SCENIC+ uses the snRNA-seq, the cisTopic object and the Topic modelling of the chromatin peaks and the motif enrichment results. For the non-multiome data (scRNA-seq and snATAC-seq not measured in the same cell) we created a pseudo-multiome by generating metacells. These metacells are created by randomly sampling 10 cells from each modality (snRNA & snATAC) and taking the average of the counts within these cells. For each of the developmental time-points, we first extracted the cell metadata and consensus regions from our ArchR objects. We then created a cisTopic object using this information along with the fragments from the raw fragment file. Next, we applied topic modeling to group the co-accessible open chromatin regions into multiple topics. We used four different quality metrics to choose the optimal number of topics. We then used pyCisTopic (version 1.0a0) to predict the candidate enhancer regions. Next, we used the wrapper of pycistarget within SCENIC+ to identify the motifs that are enriched in regions that are differentially accessible for each cell-type and are on the candidate enhancer region. For this motif enrichment we used a pre-computed motif-score database. Finally, we ran SCENIC+ to infer the GRNs.

## DATA AVAILABILITY

The single cell RNAseq, ATACseq and multiome (combined ATAC- and RNA-seq) data have been deposited in ArrayExpress; E-MTAB-14127 (snRNA-seq), E-MTAB-14128 (snATAC-seq) and E-MTAB-14129 (multiome).

## CODE AVAILABILITY

The following codes are freely available: https://github.com/galib36/developing-human-oesophagus

## ACKNOWLEDGEMENTS

We are very grateful to all women who consented to take part in our research programme and for the assistance of research nurses and clinical colleagues at the Manchester University NHS Foundation Trust. We thank Andy Hayes of the Bioinformatics and Genomic Technologies Core Facility at the University of Manchester. The work was supported by MRC Human Cell Atlas project grant MR/5036121/1. RJ was an MRC clinical research training fellow. We thank Johannes Schumacher, Rebecca Fitzgerald and Stuart MacGregor for sharing the GWAS summary statistics and also Karol Nowicki-Osuch and Rebecca Fitzgerald for sharing scRNA-seq data from BO and upper GI tissues.

## AUTHOR CONTRIBUTIONS

NAH and ADS devised the study and planned experiments. NAH, REJ, KPH were involved in study design and oversight of human embryonic material collection. AM processed the human embryonic material and prepared samples for all sequencing analyses. SMB conducted the bioinformatics analyses. CP provided input on the GWAS studies. YA provided samples and expertise on BO. ADS and NAH wrote the manuscript with input and editing from SMB and AM.

## FURTHER INFORMATION

Correspondence and requests for further information should be addressed to ADS or NAH.

---

**Supplementary Fig.1:**
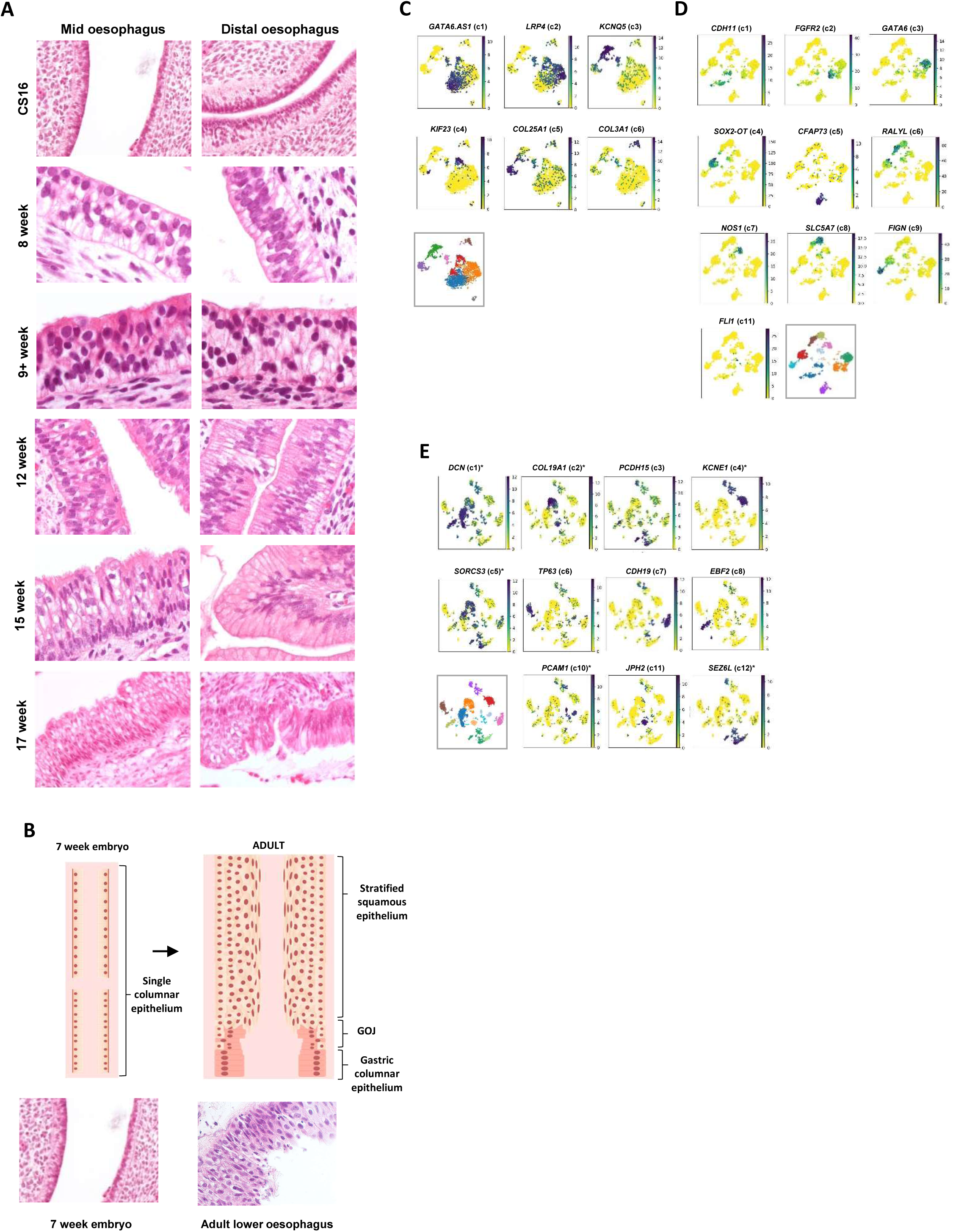
Characterisation of the developing human oesophagus. (A) Examples of H&E stained embryonic/fetal tissues at the indicated developmental stages. (B) Schematic illustration of the changing nature of the oesophageal epithelial layer in the embryo and adult. Representative H&E staining of the epithelial layers is shown below each stage. (C-E) tSNE plots with the indicated cluster marker gene expression superimposed for week 7 (C), week 9 (D) and week 15 (E). Note that week 7 clusters 7 and 8, week 9 cluster 10 and week 15 cluster 9 lack distinct marker genes which highlight the cluster. Asterisks in E indicate genes that demarcate one cluster but are also found in additional clusters.

**Supplementary Fig. 2:**
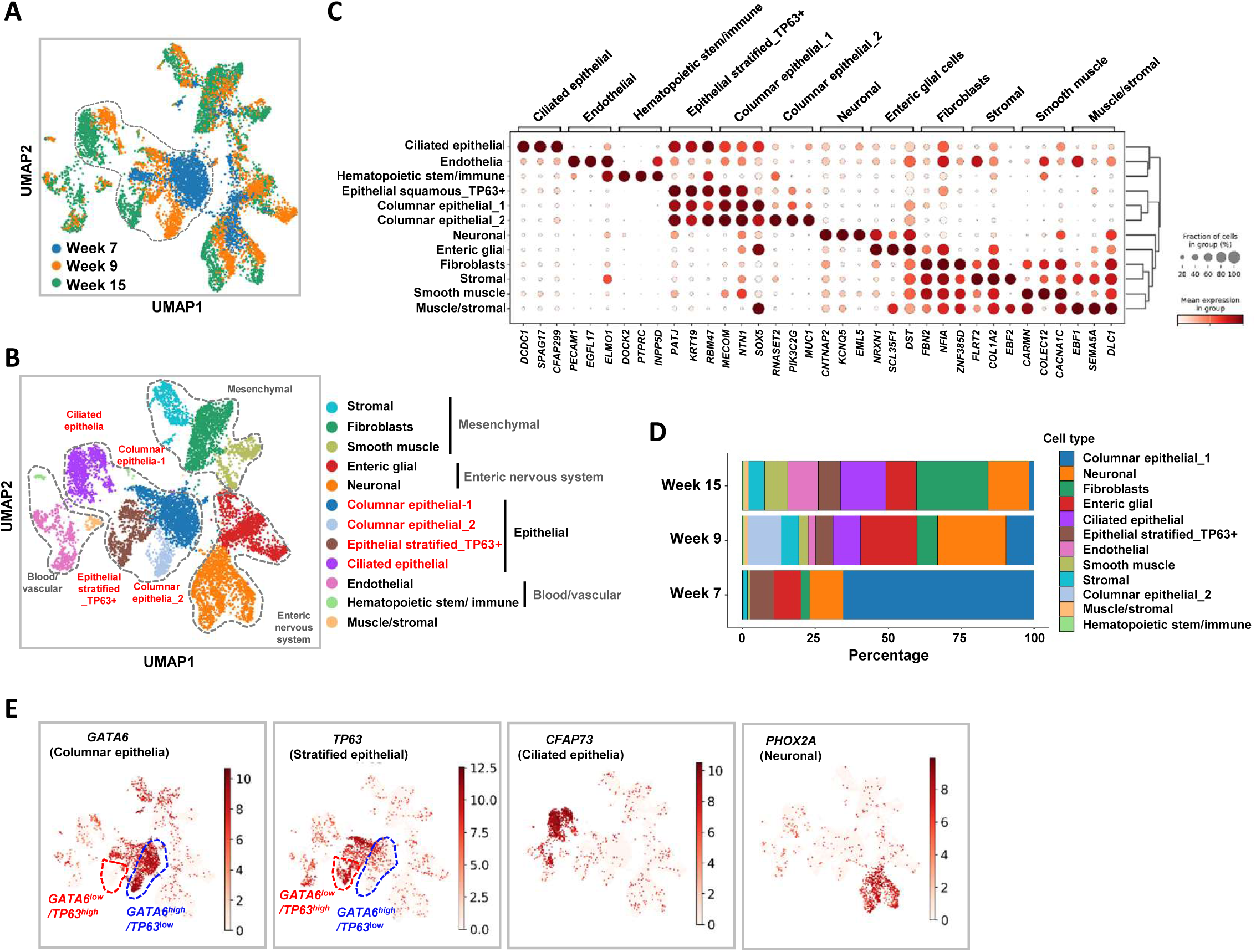
Characterisation of the cell populations in the developing oesophagus. (A and B) UMAPs of all the cell populations from the combined week 7, 9 and 15 samples. The locations of cells originating from each time point (A) and the major cell types (B) are projected on top of the clusters and grouped according to similarity. (C) Dotplot of the relative expression of three representative markers for each of the cell clusters annotated in part B. The fraction of cells expressing each marker and relative average expression levels (column normalised) are represented by the size and intensity respectively, of each dot. (D) Percentage of each of the major cell types found at each developmental timepoint. (E) Relative expression (indicated by scale bars) of the indicated genes projected on cells in the UMAP in parts A and B.

**Supplementary Fig. 3:**
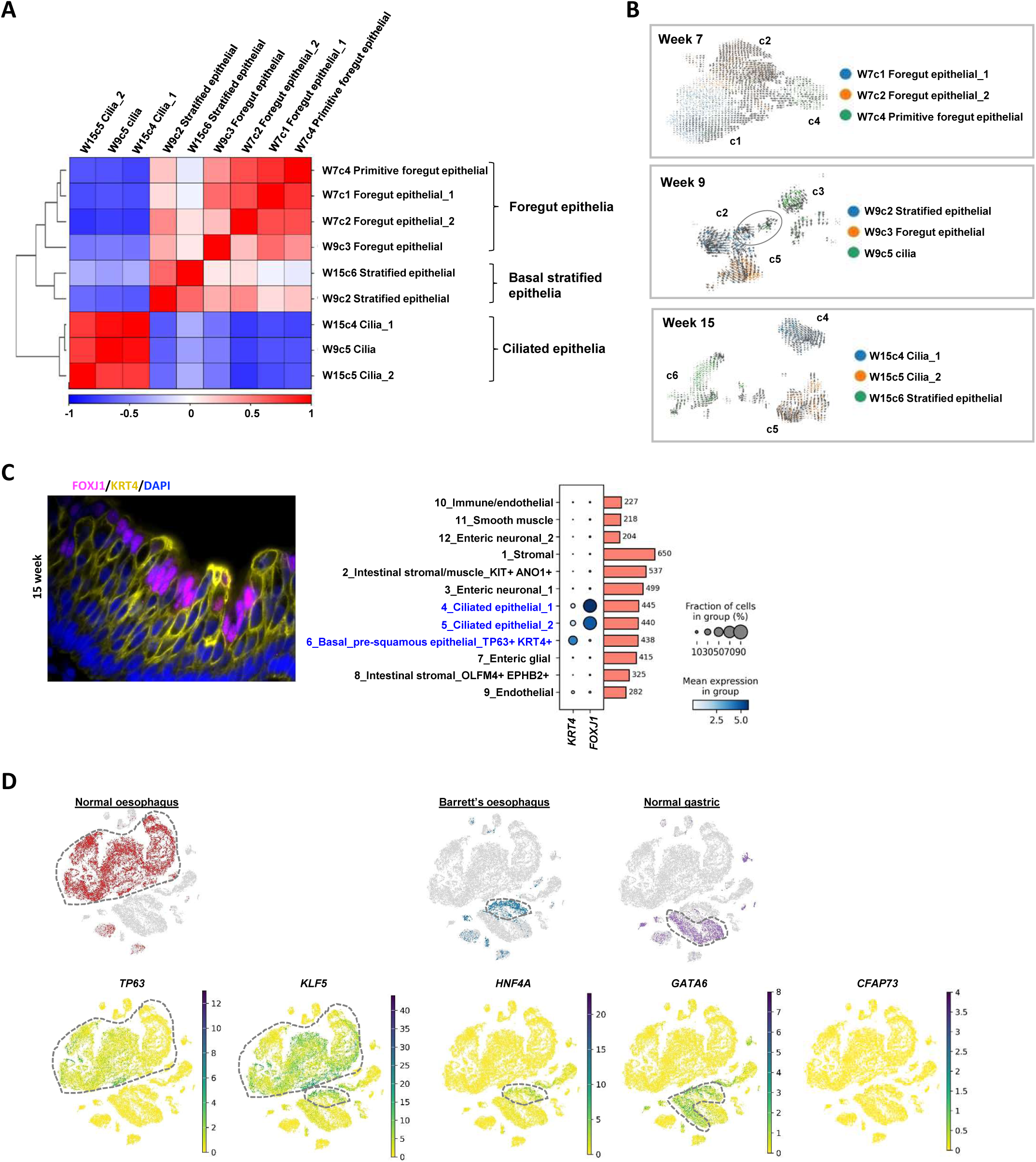
Inter relationships between epithelial cell clusters. (A) Pearson’s correlation plot comparing gene expression profiles of the epithelial cell clusters across developmental time points. Broad categories of epithelial cell types are indicated. (B) RNA velocity (scVelo) analysis of the epithelial cell clusters at each time point superimposed on tSNE plots (same plot as Fig. 2B but with more arrows). (C) Immunohistochemistry (left) and dotplots (right) showing the expression of FOXJ1 and KRT4 in the epithelial layer of 15 week embryos. (D) UMAPs of snRNA-seq data from the cell populations found in the adult upper GI tract and BO (Nowicki-Osuch et al., 2021). Cells corresponding to the different tissue or disease types are shown (top) and the expression of the indicated genes are superimposed on top of the UMAP (bottom).

**Supplementary Fig. 4:**
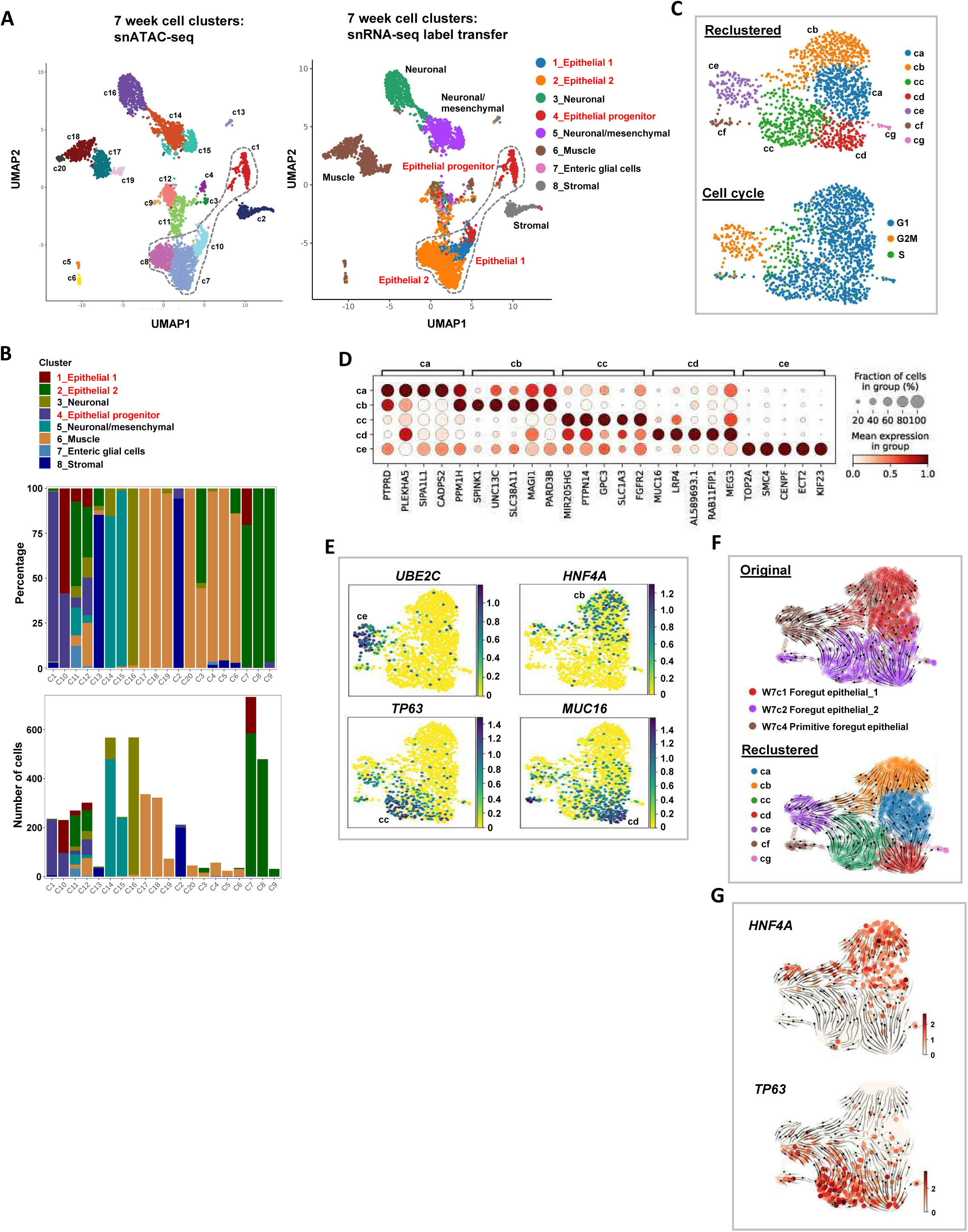
Development regulatory networks of the primitive stage epithelial cell populations. (A) UMAPs derived from snATAC-seq data showing the cell populations in 7 week embryos. Cells are clustered according to snATAC-seq signal (top), annotated based on label transfer from snRNA-seq (right). Note that the enteric glial cells mapped to cluster c11 but only represented a small proportion of the entire cluster, and along with clusters c3 and c12, cluster c11 contained numerous cell types, potentially due to technical reasons. These clusters were located close to the small clusters c9 and c4 and these five clusters could not be unambiguously defined by label transfer. (B) Proportion of cells found in each of the snATAC-seq-derived 7 week clusters matching with each snRNA-seq derived 7 week cluster after gene expression integration (top). Total numbers of cells in each cluster are also shown (bottom). (C) UMAPs derived from snRNA-seq data in 7 week embryos following reclustering into 7 clusters (top). Cell cycle stage is superimposed on this UMAP (bottom). (D) Dot plot showing the five top marker genes for clusters ca-ce. (E) UMAPs of epithelial cells in 7 week embryos with the indicated gene expression levels superimposed. (F) RNA velocity analysis showing original clusters (see Fig. 2D) (top) and reclustered epithelial cells (bottom). (G) RNA velocity analysis with expression of *HNF4A* or *TP63* superimposed on the UMAP.

**Supplementary Fig. 5:**
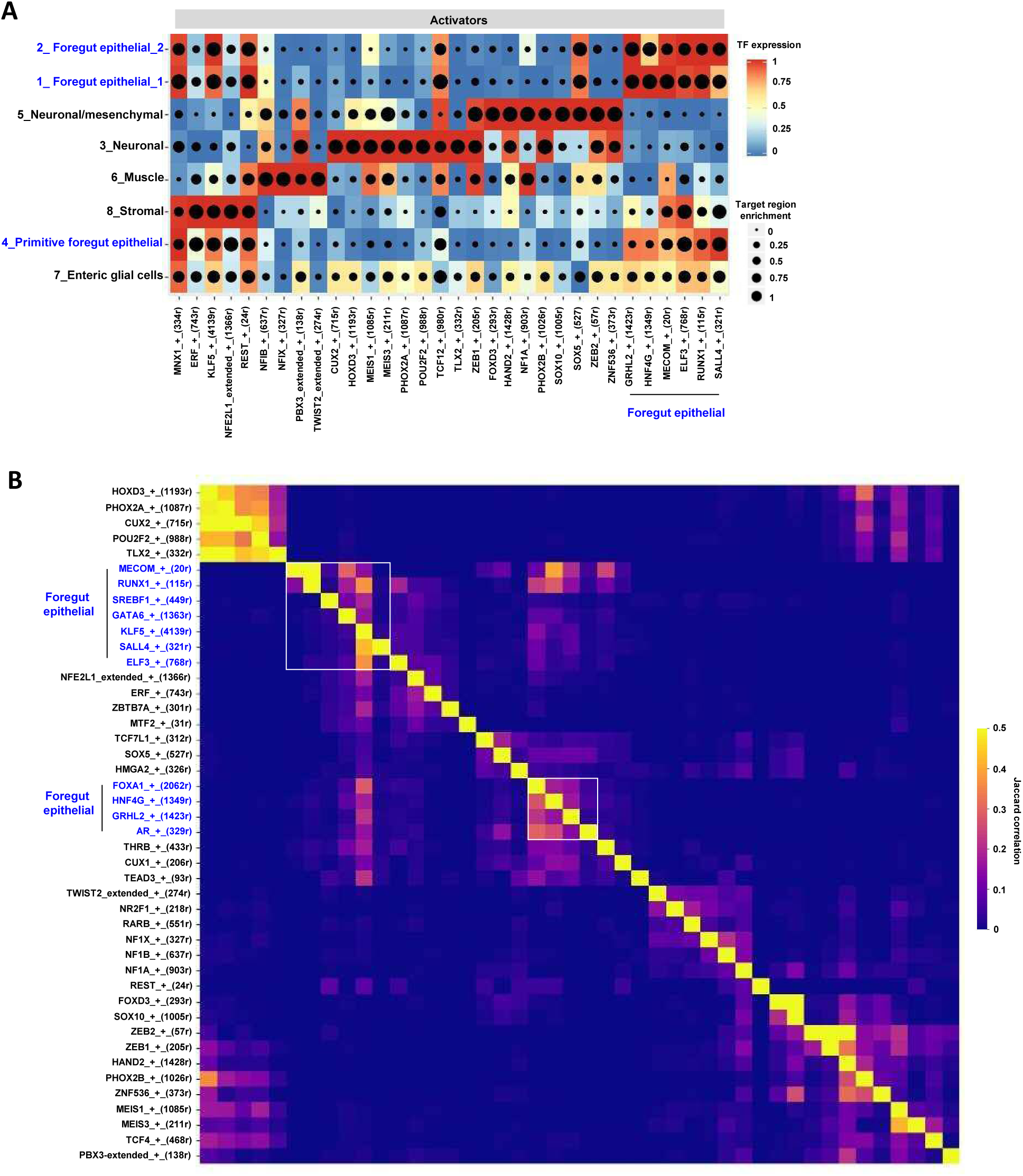
Gene regulatory networks (GRN) in the primitive stage cell populations. (A) Top scoring GRNs in each of the 7 week cell clusters (eRegulons correlation coefficient above 0.70 or below -0.75). Transcription factors and number of linked target genes (in brackets) are shown with relative TF expression levels colour coded, and target region enrichment in each cluster is shown by the size of dots. (B) Heatmap depicting an extended number of GRNs found in the 7 week cell clusters. Jaccard correlation coefficients between the different GRNs are depicted with foregut epithelial GRNs boxed.

**Supplementary Fig. 6:**
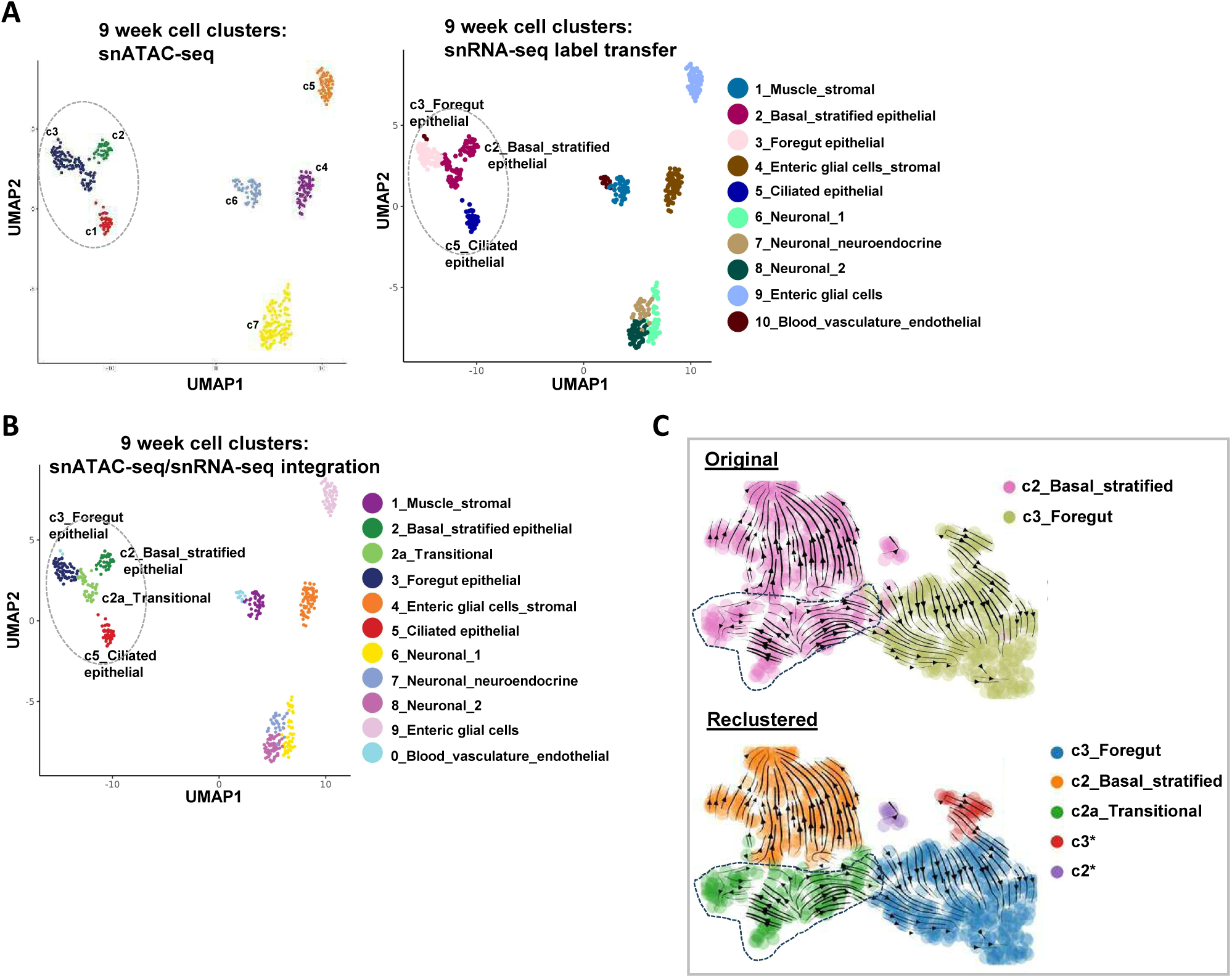
Development regulatory networks in the columnar to squamous epithelial transition. (A) UMAPs derived from snATAC-seq data showing the cell populations in 9 week embryos. Cells are clustered according to snATAC-seq signal (left), annotated based on label transfer from snRNA-seq (right). (B) UMAPs derived from snATAC-seq data from the cell populations in 9 week embryos, reclustered after label transfer from snRNA-seq. The new transitional population c2a is added to the epithelial cell clusters. Epithelial cell clusters are enclosed within the by dotted ellipses. (C) RNA velocity analysis showing original c2 and c3 epithelial cells clusters (top) and 5 new clusters when these cells are reclustered (bottom). The transitional cluster c2a is highlighted.

**Supplementary Fig. 7.**
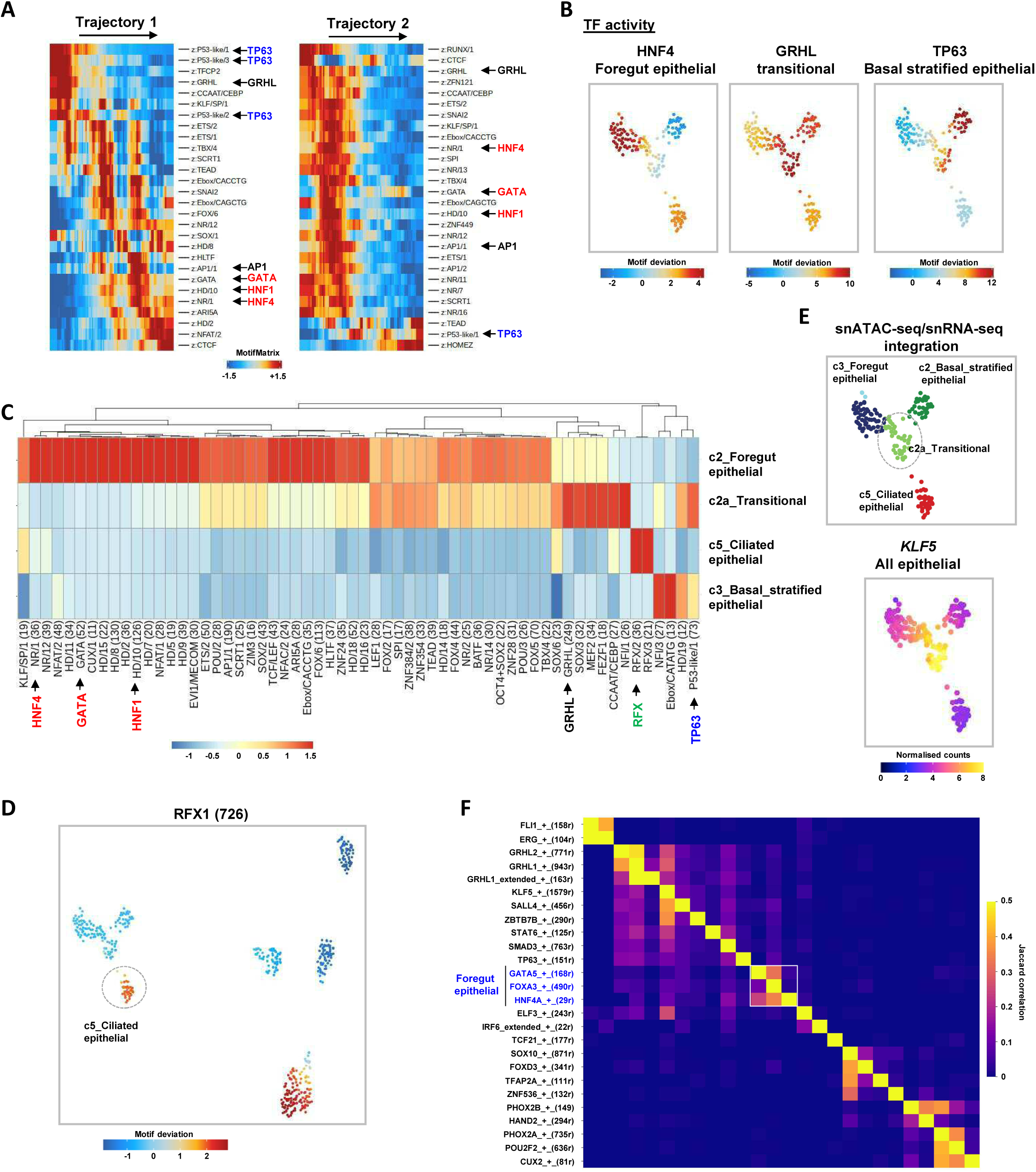
Regulatory networks in week 9 epithelial cells. (A) Motif deviation score across pseudotime with max and min value limited to 1.5 and -1.5 respectively, showing all motif labels across the trajectories depicted in Fig. 4C and D. (B) Transcription factor binding motif deviation scores for HNF4, GRHL and TP63 in individual cells projected on the epithelial population snATAC-seq-derived UMAP. (C) Heatmap showing the relative enrichment of the indicated transcription factor binding motifs in each of the epithelial cell clusters (scale bar shows scaled hypergeometric enrichment of a peak annotation). Motifs discussed in the text are highlighted. (D) Transcription factor binding motif deviation scores for RFX transcription factors in individual cells projected on all of the cell clusters in the snATAC-seq-derived UMAP. (E) UMAP of the epithelial cell clusters derived from snATAC-seq (left) showing the expression (gene integration scores from snRNA-seq) of *KLF5* (right). (F) Heatmap depicting an extended number of GRNs found in the 9 week cell clusters. Jaccard correlation coefficients between the different GRNs are depicted with foregut epithelial GRNs

**Supplementary Fig. 8:**
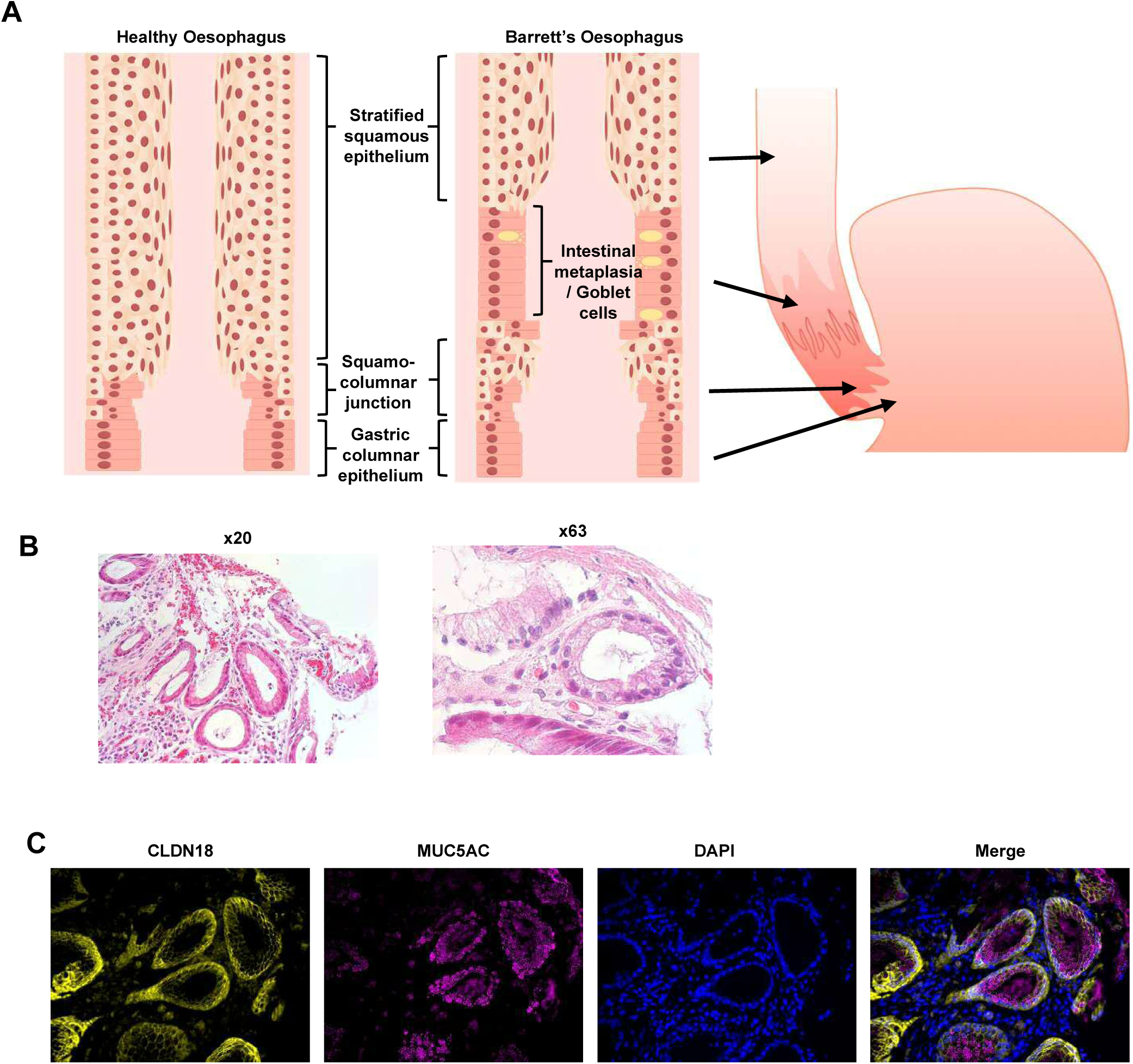
Barrett’s oesophagus tissue architecture characterisation. (A) Schematic views of the cellular organisation of adult oesophagus in healthy and Barrett’s patients. (B) H&E staining of Barrett’s sample used to generate snATAC-seq at 20x (left) and 63x (right) magnification. (C) ISH analysis of the Barrett’s marker proteins CLDN18 and MUC5AC in Barrett’s samples. Nuclei are shown by DAPI staining.

**Supplementary Fig. 9:**
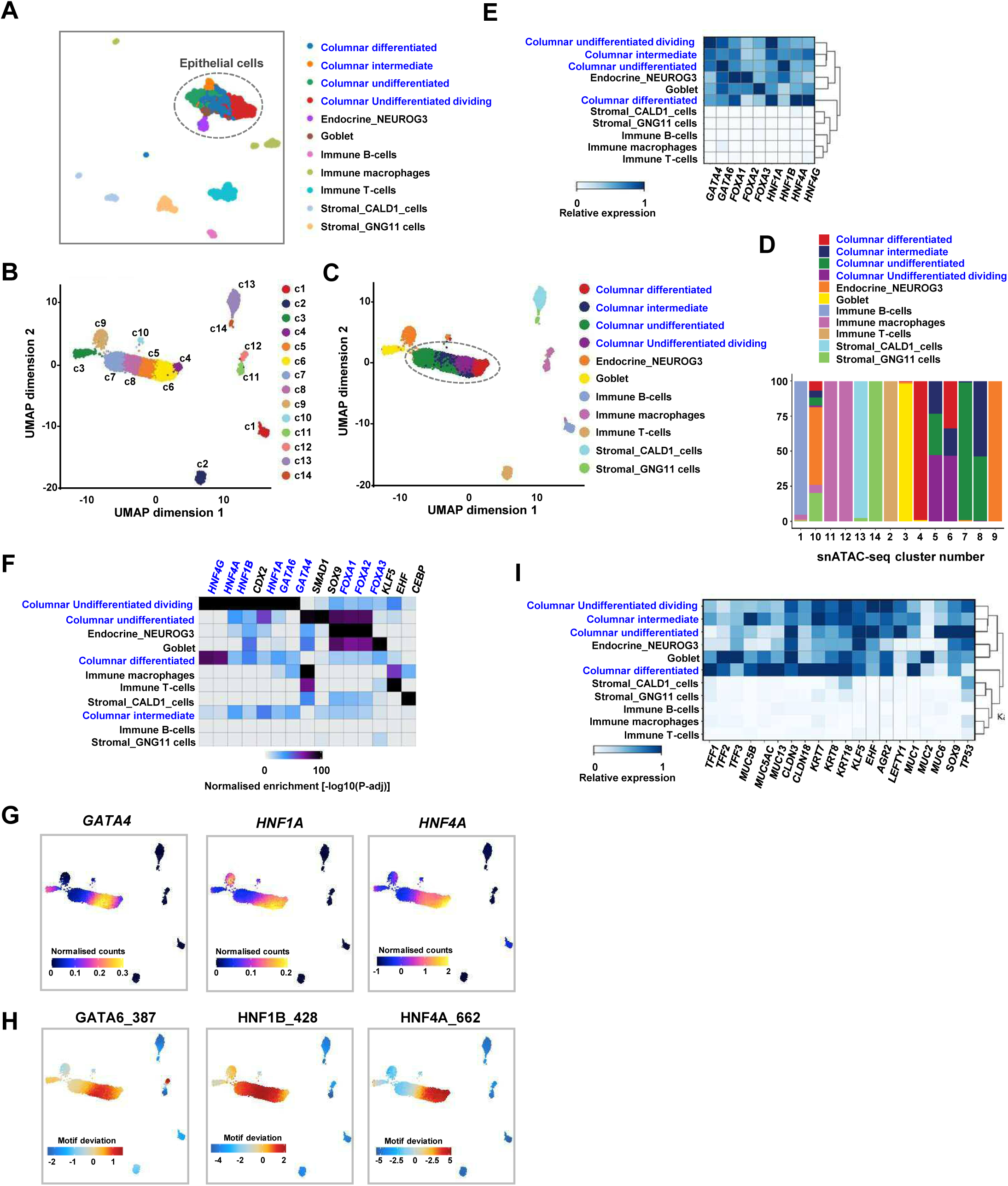
Single cell RNA-seq and ATAC-seq analysis of Barrett’s oesophagus. (A) UMAP of scRNA-seq data from a patient with non-dysplastic Barrett’s. Clusters are annotated as previously (Nowicki-Osuch et al., 2021). Epithelial cell clusters are circled. (B and C) UMAP of snATAC-seq data from a patient with non-dysplastic Barrett’s. Unsupervised clustering is shown (B) and the cell clusters are given identities by label transfer from the scRNA-seq clusters (C). Columnar epithelial cell clusters are circled. (D) Contributions of cells identified by label transfer to each of the ATAC-seq clusters. (E) Heatmap showing the relative expression of the indicated transcription factors in each of the Barrett’s sample clusters. (F) Heatmap showing the relative enrichment of DNA binding motifs for each of the indicated transcription factors in each of the Barrett’s sample clusters. Enrichment values (-log10(p-adj)) are scaled between 0 to 100. (G) Gene expression scores (from gene integration matrix) for the indicated transcription factors for individual cells projected on the epithelial population ATAC-seq-derived UMAP. (H) Transcription factor binding motif scores for individual cells projected on the epithelial population ATAC-seq-derived UMAP. (I) Heatmap showing the relative expression of the indicated Barrett’s marker genes (scaled for each column) in each of the Barrett’s sample scRNA-seq derived clusters.

**Supplementary Fig. 10.**
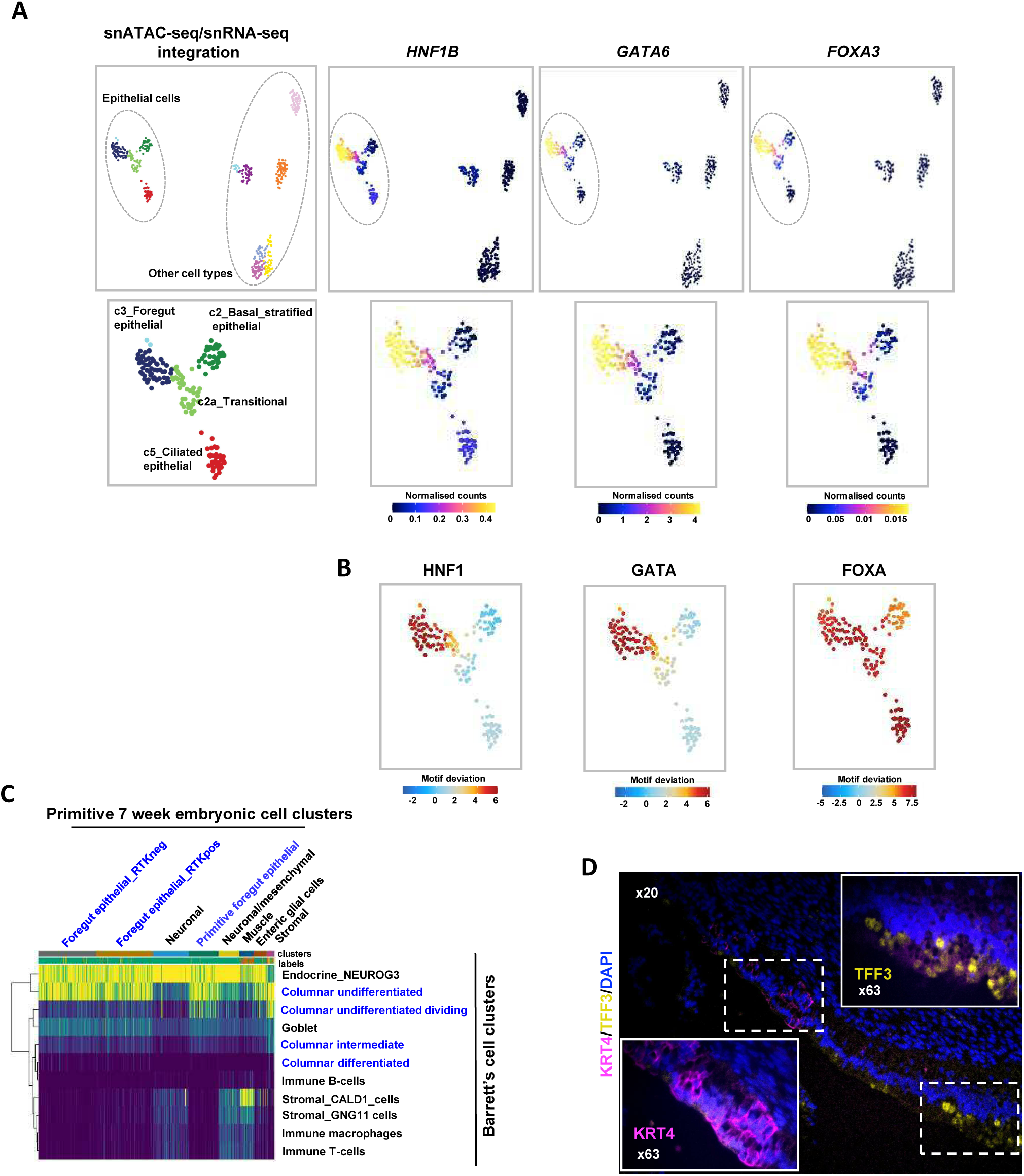
Barrett’s oesophagus resembles early developmental cell states. (A) UMAPs of week 9 developmental clusters derived from snATAC-seq (see Fig. 4) showing cluster identifications (leftmost panels) or with the expression of the indicated transcription factors (by gene integration from RNAseq) projected on top of these (right panels). Epithelial cell populations are highlighted below the complete heatmaps. (B) Transcription factor motif scores in each cell projected on top of the epithelial cell clusters. (C) Heatmap showing similarity scores between each cell in the 7 week developmental clusters (x-axis) and the corresponding cell types found in Barrett’s cell clusters (y-axis). (D) Immunofluorescence staining of 9 week oesophageal tissue for the nuclear marker DAPI, the squamous epithelial marker KRT4 (pink) and also TFF3 (green) which highlights secretory columnar cells in the lower oesophagus. Higher magnification insets are shown of the upper (left) and lower oesophagus (right).

**Supplementary Fig. 11.**
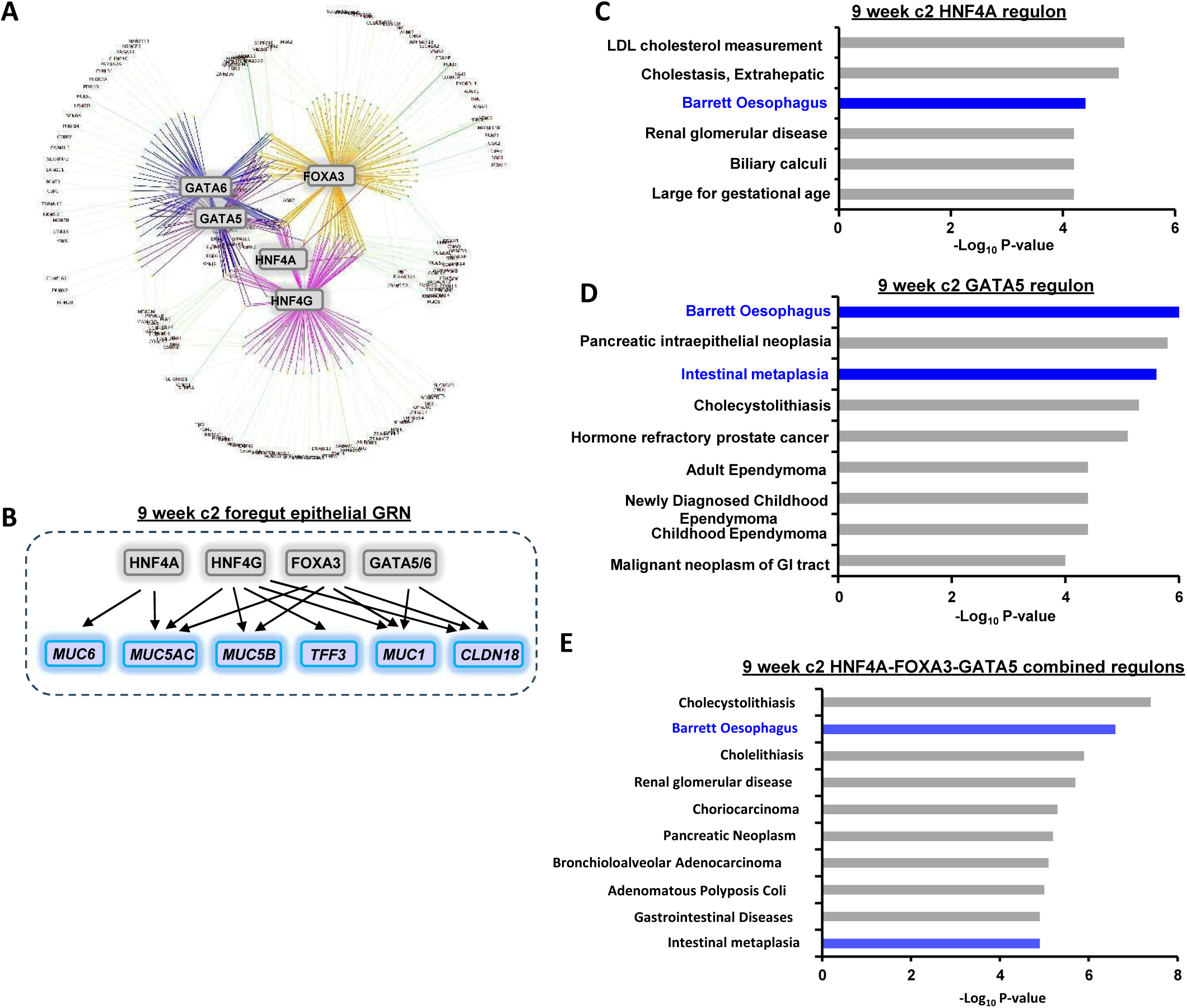
Barrett’s-like GRNs in the foregut epithelial cells of transitory state embryos. (A) Broad view of the gene regulatory networks controlled by the GATA-FOXA-HNF4 TF axis. (B) Regulatory links derived from the week 9 foregut epithelial cluster GRN, depicting upstream -FE) Enriched DisGeNET GO terms for the HNF4A regulon (29r)(D), GATA5 regulon (168r)(E) and combined HNF4A (29r), FOXA3 (490r) and GATA5 (168r) (F) in 9 week c2 foregut epithelial cells.

**Supplementary Fig. 15.**
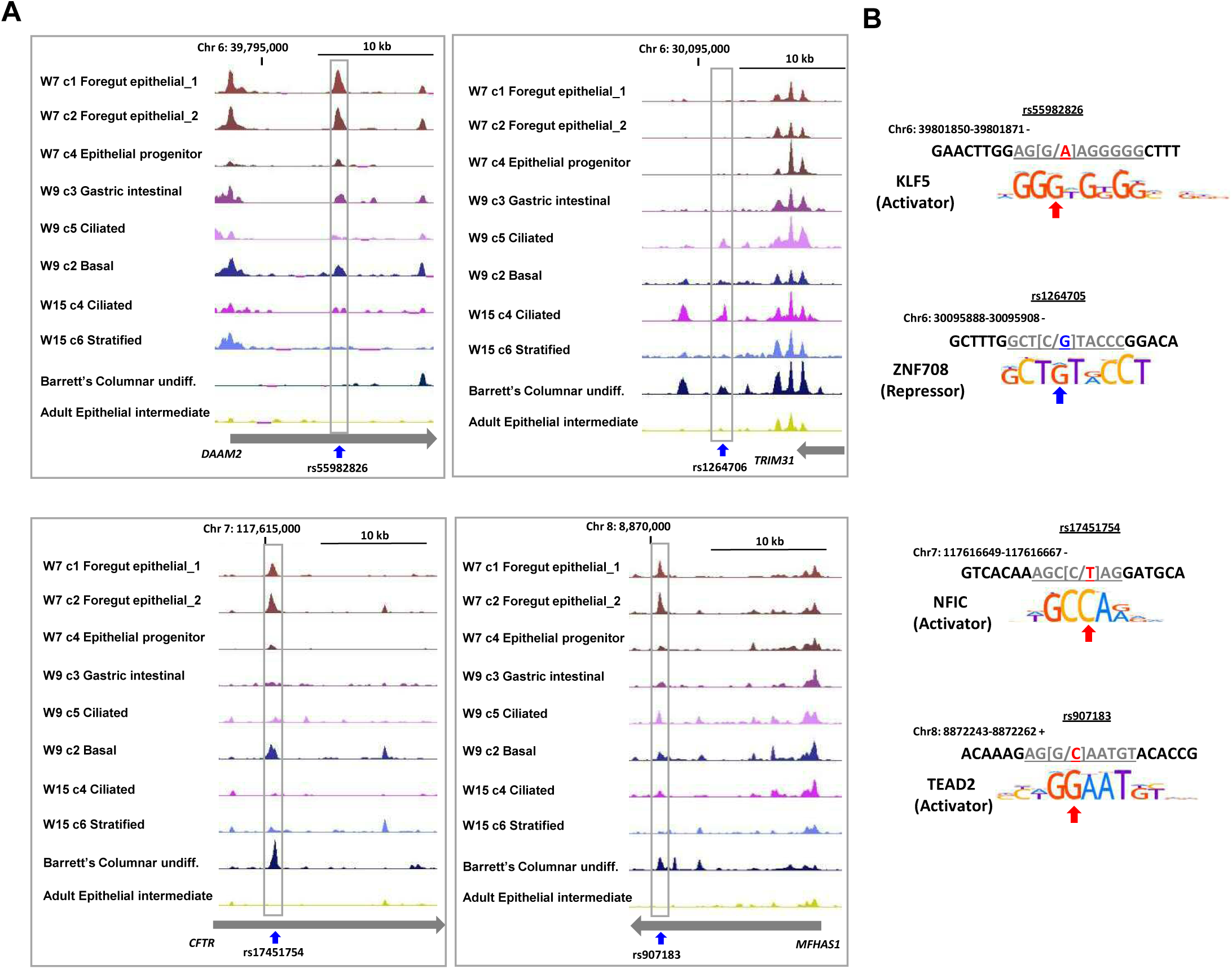
Barrett’s-associated SNPs map to enhancers in the developing oesophagus. (A) UCSC genome browser view of ATAC-seq signals surrounding SNPs rs55982826, rs1264706, rs17451754 and rs907183 in the indicated clusters of epithelial cells in human embryos, Barrett’s undifferentiated columnar epithelia and adult oesophageal epithelial cells. The arrows below the tracks represent the directionality and extent of the genes covered by the views. Peaks containing the SNPs are boxed. (B) Sequences around significantly associated GWAS SNPs. The genomic location and DNA strand is shown above the sequences. Base changes are shown as WT/risk allele and the risk allele is coloured as red for disruption of an activator binding site or blue for creation of a repressor binding site. Logos for the DNA binding motifs of the indicated transcription factors are shown and corresponding bases in the DNA sequence are underlined. Arrows indicate the bases changed by the risk SNP.

**Supplementary Fig. 12:**
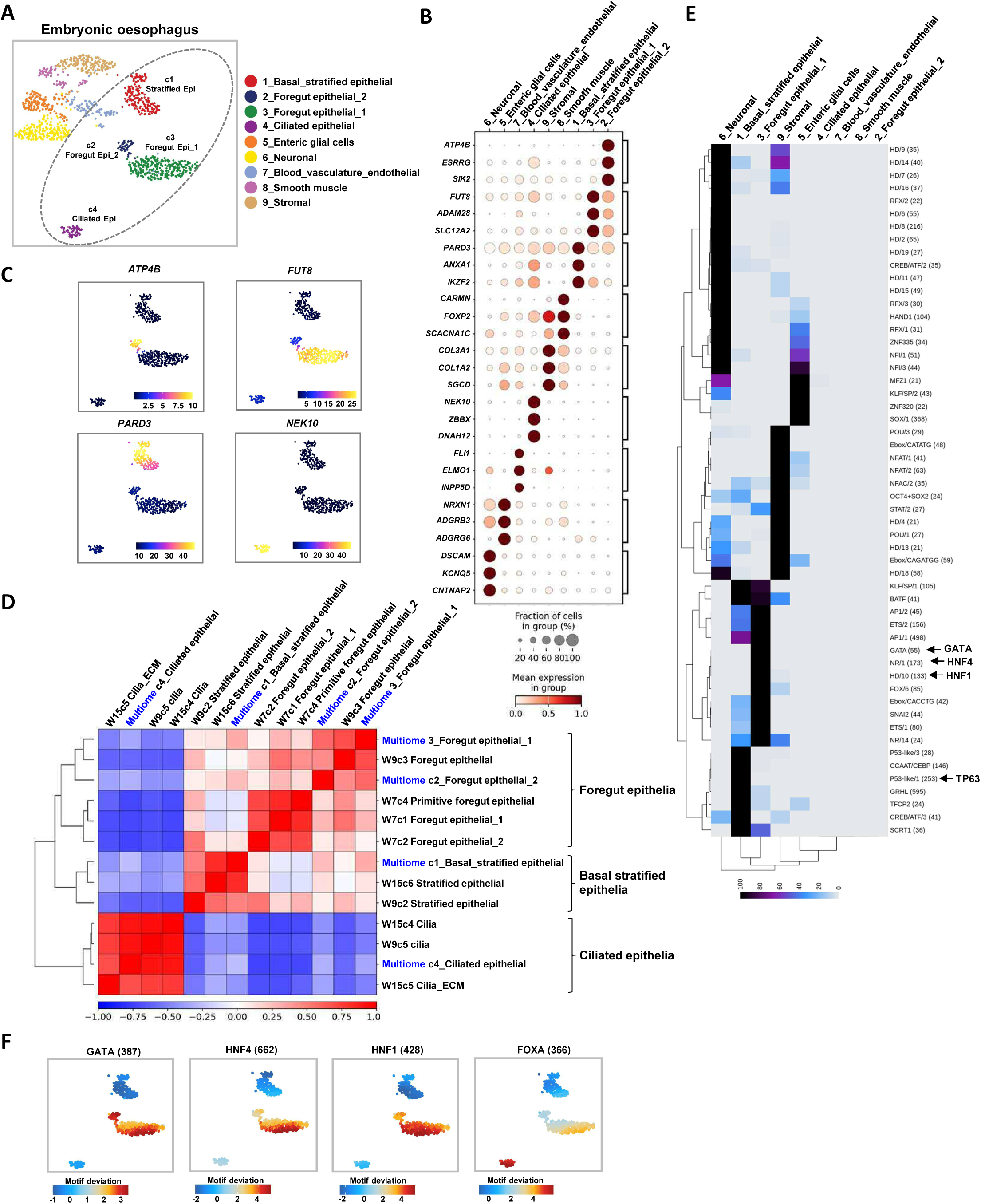
Multiome analysis of the developing oesophagus at the embryonic-fetal transition. (A) UMAP of embryonic multiome cell clusters based on joint embedding of snATAC-seq and snRNA-seq. Major epithelial populations are annotated on the plot. (B) Dotplot of the relative average expression of three representative markers for each of the cell clusters found in the embryonic multiome data. The fraction of cells expressing each marker and relative expression levels (column normalised) are represented by the size and intensity respectively, of each dot. (C) Relative expression (indicated by scale bars) of the indicated genes projected on cells in the UMAP in part A. (D) Pearson’s correlation plot comparing gene expression profiles of the epithelial cell clusters across developmental time points from Week 7, 9 and 15 snRNA-seq and embryonic multiome data. Broad categories of epithelial cell types are indicated. (E) Heatmap showing the relative enrichment of the indicated transcription factor binding motifs in each of the multiome cell clusters. Motifs discussed in the text are highlighted. (F) Transcription factor binding motif deviation scores for individual cells projected on the epithelial populations in the UMAP in part A.

**Supplementary Fig. 13:**
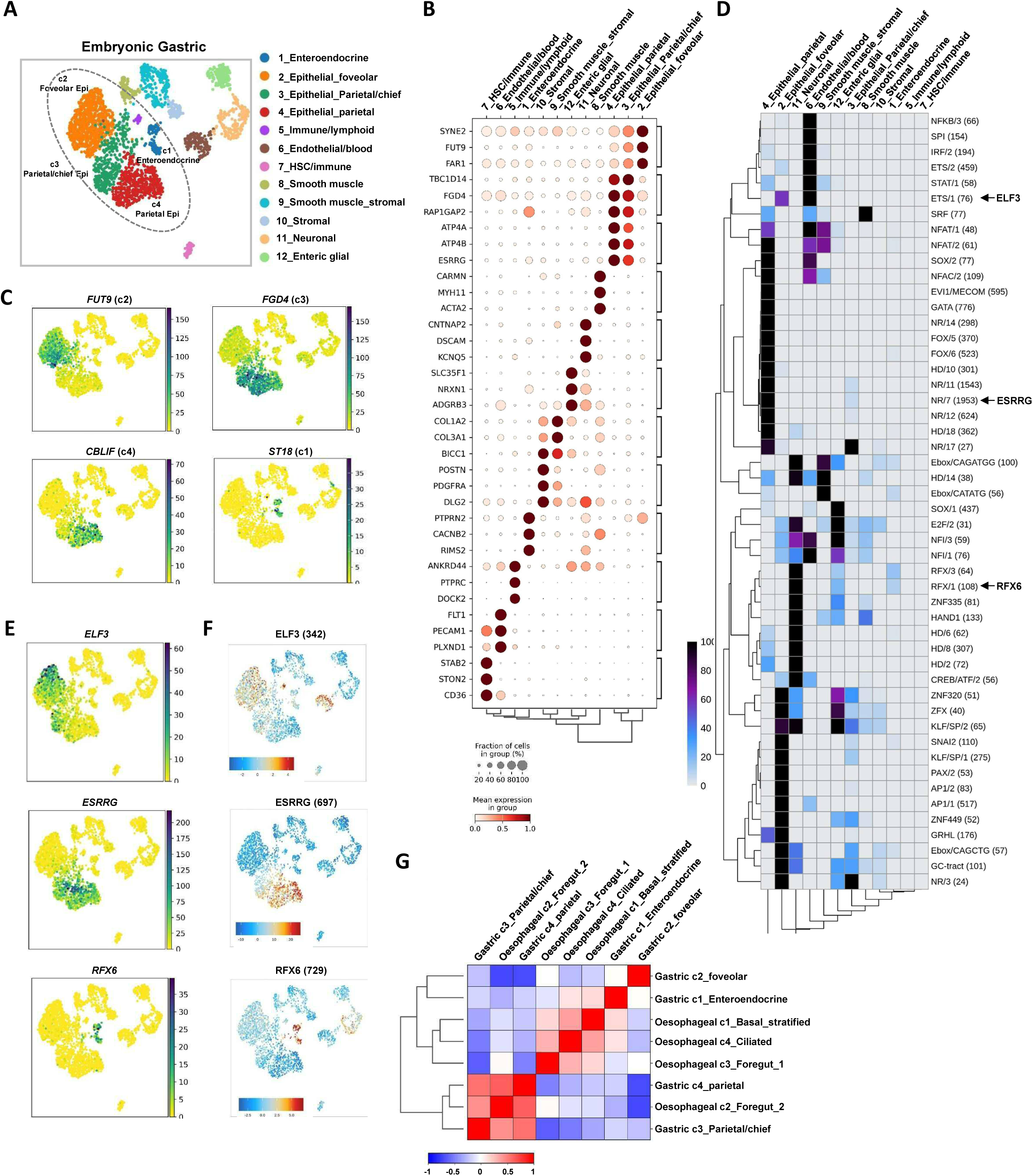
Multiome analysis of the developing gastric cardia at the embryonic-fetal transition. (A) UMAP of embryonic multiome cell clusters based on joint embedding of snATAC-seq and snRNA-seq. Major epithelial populations are annotated on the plot. (B) Dotplot of the relative average expression of three representative markers for each of the cell clusters found in the embryonic multiome data. The fraction of cells expressing each marker and relative expression levels (column normalised) are represented by the size and intensity respectively, of each dot. (C) Relative expression (indicated by scale bars) of the indicated genes projected on cell clusters in the UMAP in part A. (D) Heatmap showing the relative enrichment of the indicated transcription factor binding motifs in each of the multiome cell clusters. Motifs discussed in the text are highlighted. (E) Relative expression (indicated by scale bars) of the indicated transcription factor encoding genes projected on cell clusters in the UMAP in part A. (F) Transcription factor binding motif deviation scores for individual cells projected on the epithelial populations in the UMAP in part A. (G) Pearson’s correlation plot comparing gene expression profiles of the epithelial cell clusters from the embryonic oesophageal and gastric multiome data. Broad categories of epithelial cell types are indicated.

**Supplementary Fig. 14:**
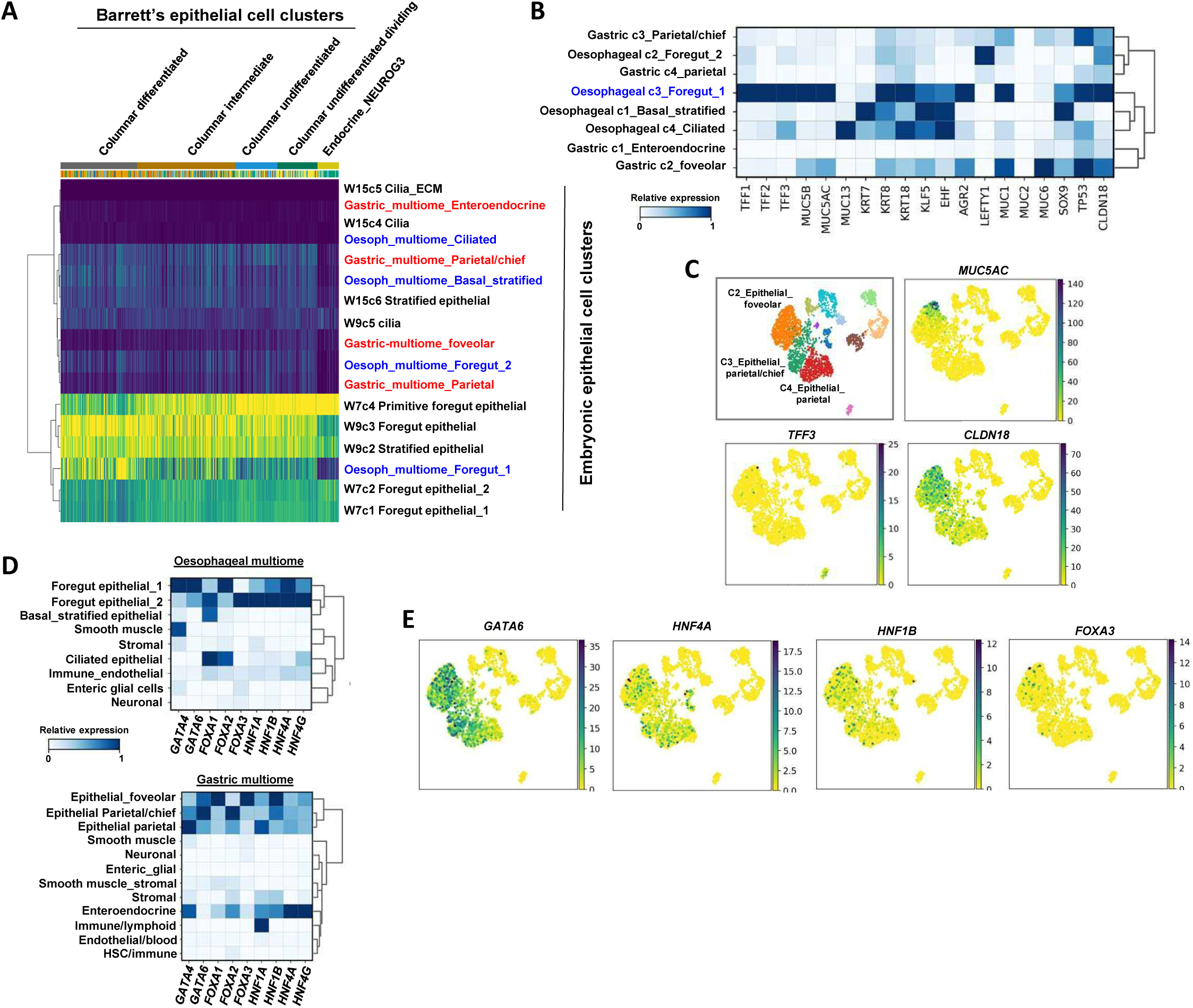
Comparison of the developing gastric and oesophageal epithelial populations with Barrett’s oesophagus. (A) Heatmap showing similarity scores between each cell in the epithelial cell clusters from adult Barrett’s (x-axis) and the corresponding cell types found in different embryonic epithelial cell clusters (y-axis). (B and D) Heatmaps showing the relative expression (column normalised) of the indicated Barrett’s marker genes (B) or the indicated transcription factors (D) in all epithelial clusters from the gastric and oesophageal multiomes (B) or each of the cell clusters from the embryonic oesophageal multiome data (D, top) or the embryonic gastric multiome (D, bottom). (C and E) UMAPs of the epithelial cell clusters derived from gastric multiome (C, top left) showing the expression of the indicated Barrett’s marker genes (C) or Barrett’s core regulatory transcription factors (E). Major gastric epithelial cell clusters (c2-4) are labelled.

