## Supplementary Table S4 for "The metaplastic precursor state to oesophageal adenocarcinoma represents reversion to a transient epithelial cell state in the developing oesophagus"

**Supplementary Table S4**. Antibodies used in this study.

| Antigen | Manufacturer | Ref Number | Dilution | Raised in |
| --- | --- | --- | --- | --- |
| CLDN18 | Abcam | ab203563 | 1:500 | Rabbit |
| FOXJ1 | Abcam | ab235445 | 1:1000 | Rabbit |
| KRT4 | Antibodies.com | A249123 | 1:500 | Mouse |
| MUC5AC | Cell Signalling | 61193T | 1:400 | Rabbit |
| MUC5AC | Novus | NBP2-15196 | 1:500-2000 | Mouse |
| TFF3 | Abcam | ab109104 | 1:500-2000 | Rabbit |
